# Composition can buffer protein dynamics within liquid-like condensates

**DOI:** 10.1101/2022.05.16.492059

**Authors:** Stela Jelenic, Janos Bindics, Philipp Czermak, Balashankar R Pillai, Martine Ruer, Carsten Hoege, Alex S Holehouse, Shambaditya Saha

## Abstract

Most non-membrane-bound compartments in cells that form via phase separation have complex composition. While phase separation of individual proteins that form these compartments is well-documented, the mechanisms that modulate dynamics of individual proteins in multicomponent systems remain unclear. Here, we used in vitro reconstitution and in vivo experiments to investigate how the dynamics of a scaffold protein PGL-3 is regulated within the liquid-like ‘P granule’ compartment in *C. elegans*. Using mutational and biophysical perturbations, we generated PGL-3 constructs that form condensates in vitro with widely varying dynamics. Using these PGL-3 constructs, we show that introducing other P granule components buffers against change of dynamics within liquid-like condensates. This dynamics-buffering effect is mediated by weak interactions among two or more components. Such dynamics-buffering may contribute to robust functional output of cellular liquid-like compartments.

## Introduction

To organize biochemistry, cells assemble compartments with distinct compositions. Many of these compartments are not enclosed within lipid membranes and assemble via liquid-liquid phase separation (LLPS) of proteins and RNAs from the surrounding cytoplasm or nucleoplasm (Banani *et al*, 2017; Brangwynne *et al*, 2009; Lyon *et al*, 2021). Studies over the past decade have focused on identifying the molecular mechanisms of LLPS within cells. The current model is that LLPS is driven by ‘scaffold’ proteins and/or RNAs which participate in homotypic or heterotypic interactions with each other to assemble cellular non-membrane-bound compartments (Van Treeck & Parker, 2018; Banani *et al*, 2016, 2017; Lyon *et al*, 2021; Riback *et al*, 2020; Sanders *et al*, 2020; Saha *et al*, 2016). Other macromolecules called ‘clients’ do not contribute to compartment assembly but concentrate within those compartments and modulate biological function. It is now appreciated that LLPS can be driven by interactions of different types i.e. electrostatic, polar or hydrophobic interactions involving both folded domains and intrinsically disordered regions (Banani *et al*, 2017; Lyon *et al*, 2021). The mechanisms that regulate dynamics of components within these compartments, however, remain unclear.

Macromolecules can phase separate into condensates with different material properties, ranging from liquid-like phases with fast dynamics to dynamically-arrested gels as well as glasses (Banani *et al*, 2017; Jawerth *et al*, 2020). Many liquid-like condensates *in vitro* have also been shown to mature over time, starting as liquid-like droplets that transition into non-dynamic assemblies (Banani *et al*, 2017). The rates of biochemical reactions within biomolecular condensates must depend on the diffusion times of the components. Recent *in vitro* investigations of LLPS for simple two-component systems have shown that the behavior of one component can modulate the dynamics of another within condensates (Boeynaems *et al*, 2019; Rhine *et al*, 2020; Zhang *et al*, 2015; Alshareedah *et al*, 2021; Boyko *et al*, 2020). Cellular condensates contain hundreds of different proteins and RNAs (e.g. P granules, P bodies and stress granules (Youn *et al*, 2019; Phillips & Updike, 2022)). How such complex composition affects dynamics of macromolecules in liquid-like compartments remains poorly understood.

P granules are a canonical liquid-like compartment found in the germline of *C. elegans* (Strome & Wood, 1982; Brangwynne *et al*, 2009). mRNA and over 70 proteins are known to concentrate in P granules (Phillips & Updike, 2022). Mutations in P granule proteins are associated with loss of fertility and fecundity in *C. elegans* (Phillips & Updike, 2022; Updike & Strome, 2010). Eight proteins – PGL-1, PGL-3, GLH-1, GLH-2, GLH-3, GLH-4, DEPS-1 and MIP-1 are currently known to be constitutive P granule components i.e., these proteins concentrate within P granules at all stages of germline development (Kawasaki *et al*, 1998, 2004; Spike *et al*, 2008a, 2008b; Price *et al*, 2021; Cipriani *et al*, 2021; Phillips & Updike, 2022). Proteins of the PGL family are specific to the *Caenorhabditis* genus and contain RNA-binding motifs. The GLH-proteins belong to the highly conserved family of DEAD-box RNA helicases. Based on current estimates of the concentrations of these proteins in vivo, the PGL- and GLH-proteins are among the most abundant constitutive components in P granules (Saha *et al*, 2016). In vitro reconstitution experiments have shown that, among the constitutive P granule proteins, only the protein PGL-3 can phase separate into liquid-like condensates in vitro (Saha *et al*, 2016, 201; Hanazawa *et al*, 2011). Two other proteins that concentrate in P granules – MEG-3 and LAF-1 can also phase separate into liquid-like condensates in vitro (Smith *et al*, 2016; Elbaum-Garfinkle *et al*, 2015). This suggests that phase separation of these proteins also contribute to the assembly of P granules at some stages of germline development. The interactions that support phase separation of the PGL-3 protein remain unclear. Structural studies on the orthologous PGL-1 protein revealed two dimerization domains (Aoki *et al*, 2016, 2021). This raises the possibility that PGL-3 also contains similar dimerization domains, and phase separation depends on interactions involving these domains. The PGL-3 protein can phase separate in vitro only at high ‘non-physiological’ protein concentrations (Saha *et al*, 2016). At physiological concentrations (approx. 1 μM), binding to mRNA facilitates phase separation of the PGL-3 protein (Saha *et al*, 2016). It remains unclear if interactions among the dimerization domains cooperate with mRNA binding to support phase separation of the PGL-3 protein.

Here, we use in vitro reconstitution and in vivo experiments in adult *C. elegans* gonads to investigate how complex composition within the P granule compartment contributes to the dynamics of one of its scaffold proteins PGL-3 (Saha *et al*, 2016; Hanazawa *et al*, 2011). We begin with structure-function analysis for insights into the mechanism of phase separation of the PGL-3 protein. Using a combination of mutational analysis and biophysical perturbations, we generated PGL-3 constructs that phase separate in vitro into condensates with widely varying dynamics. Investigating the effect of other P granule components on phase separation of PGL-3 constructs, we uncovered a role of complex composition in buffering against large change of dynamics within cellular liquid-like compartments. This “dynamics-buffering” effect is distinct from active processes that depend on ATP hydrolysis, but is mediated by weak interactions among components in liquid-like compartments.

## Results

### Homotypic and heterotypic interactions drive phase separation of PGL-3 and mRNA at physiological concentrations

We began by investigating the role of different domains in the phase separation properties of the protein PGL-3 in vitro (Supplementary Fig. 1). Together with other proteins and mRNA, PGL-3 drives the assembly of the canonical liquid-like P granules in the germline of *C. elegans*. Bioinformatic analysis of the 693 residue-long PGL-3 protein predicted that the N-terminus accounting for ~65% of PGL-3 is predominantly α-helical, while the C-terminus accounting for ~35% is mostly disordered (Fig. 1a, Supplementary Fig. 2a, 2b). Consistent with the bioinformatics predictions, circular dichroism (CD) spectroscopy under conditions that do not support phase separation showed that the PGL-3 N-terminal fragment 1-452 is predominantly α-helical, while the C-terminal fragment 370-693 is disordered (Fig. 1f).

**Fig. 1.**
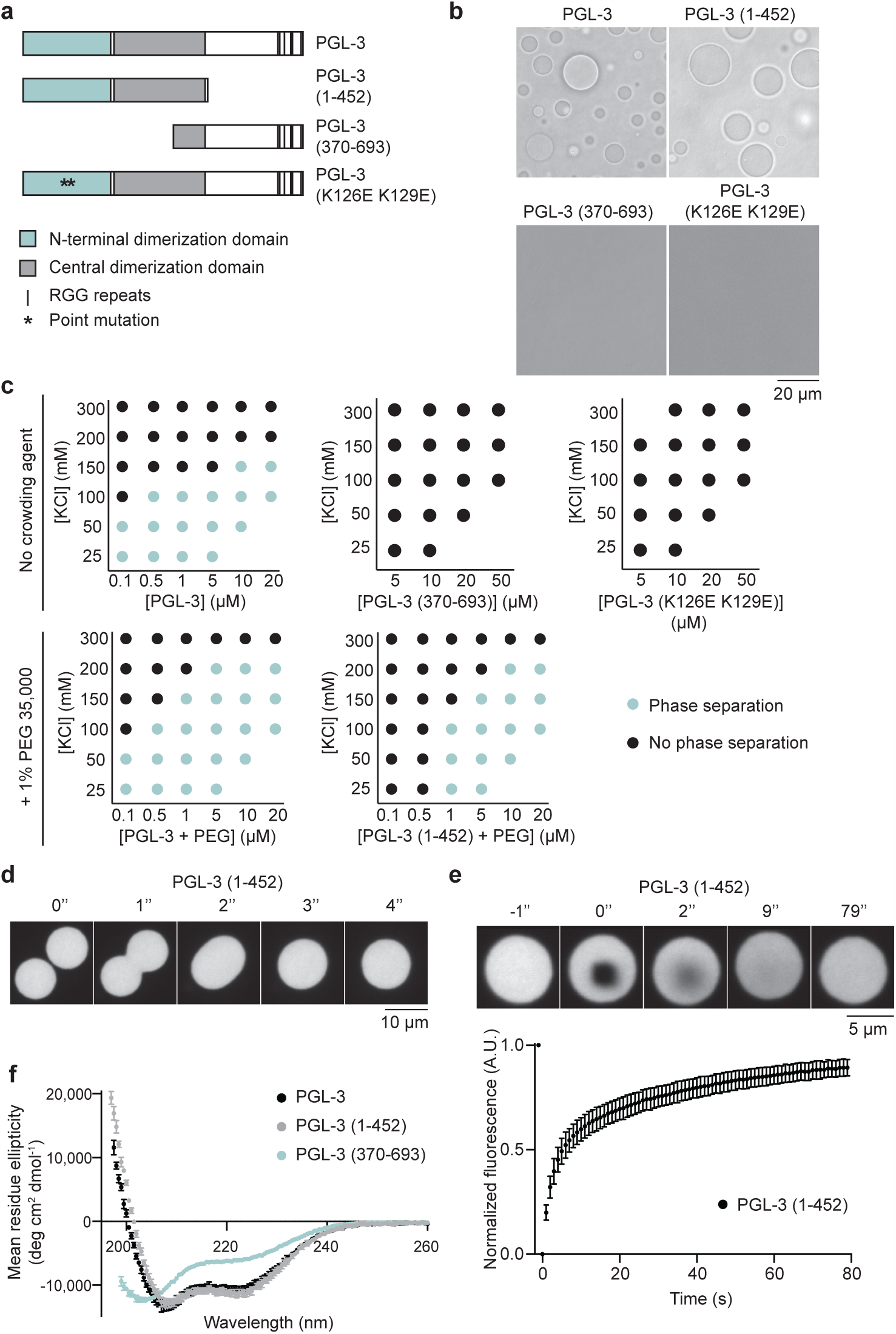
α-helical N terminus drives phase separation of PGL-3. **a** Cartoon representing PGL-3 and mutants. **b** Bright field micrographs of PGL-3 (23 μM), PGL-3 (1-452) (22 μM), PGL-3 (370-693) (67 μM) and PGL-3 (K126E K129E) (108 μM) in 25 mM HEPES pH 7.5, 100 mM KCl, 1 mM DTT in absence of crowding agents. Scale bar 20 μm. **c** Phase diagrams of PGL-3 and mutants at different protein and salt (KCl) concentrations. Blue circles represent conditions where phase separation was observed. Black circles represent conditions where phase separation was absent. Top row: Phase diagrams of PGL-3, PGL-3 (370-693), and PGL-3 (K126E K129E) in 25 mM HEPES pH 7.5, 1 mM DTT in absence of crowding agents. 5% of the pool of protein was tagged with mEGFP in each case. Bottom row: Phase diagrams of PGL-3, and PGL-3 (1-452) in 25 mM HEPES pH 7.5, 1 mM DTT, 1% PEG 35000. 5% of the pool of protein was tagged with mEGFP in each case. **d** Time-lapse fluorescence micrographs show fusion of two PGL-3 (1-452) condensates (5% tagged with mEGFP in 25 mM HEPES pH 7.5, 100 mM KCl, 1 mM DTT, 1% PEG 35000). Scale bar 10 μm. **e** Time-lapse fluorescence micrographs show fluorescence recovery after photobleaching in a condensate of PGL-3 (1-452) (5% tagged with mEGFP in 25 mM HEPES pH 7.5, 100 mM KCl, 1 mM DTT, 1% PEG 35000). Scale bar 5 μm. Shown below is a plot of average normalized fluorescence in bleached regions (1.5 μm × 1.5 μm) of the PGL-3 (1-452) condensates (n = 24) over time. Error bars represent 1 SD. **f** Plot of mean residue ellipticity of PGL-3 and mutants vs. wavelength assayed using circular dichroism (CD) spectroscopy. Black circles: PGL-3 (3 μM, 10 mM Potassium phosphate pH 7.5, 300 mM KCl, 1 mM TCEP), Grey circles: PGL-3 (1-452) (3 μM, 10 mM Potassium phosphate pH 7.5, 300 mM KCl, 1 mM TCEP), Blue circles: PGL-3 (370-693) (10 μM, 10 mM Potassium phosphate pH 7.5, 150 mM KCl, 1 mM TCEP). Error bars represent 1 SD among six CD scans.

Intrinsically disordered polypeptides have in many cases been shown to be necessary and sufficient to drive phase separation. We therefore tested if a recombinantly purified C-terminal fragment of PGL-3 (370-693), can drive phase separation in vitro. To our surprise, we did not observe any phase separation even at high protein concentrations up to ~70 µM (Fig. 1b, c). This greatly exceeds the physiological concentration of the PGL-3 protein in *C. elegans* (estimated to be ~1 µM) (Saha *et al*, 2016). However, we found that a recombinantly purified N-terminal fragment of PGL-3 (1-452) phase separates into condensates at close to physiological concentrations of PGL-3 (Fig. 1b, c, Supplementary figure 2e). Similar to full-length PGL-3, condensates of the N terminal PGL-3 fragment (1-452) are liquid-like in nature. First, condensates are spherical in shape (Fig. 1d, e, Supplementary Fig. 2d). Second, condensates fuse over time to generate a larger condensate (Fig. 1d). Third, fluorescence recovery after photobleaching (FRAP) experiments showed that PGL-3 (1-452) molecules intermix rapidly (t_1/2_ = 4 s) within a condensate (Fig. 1e). We conclude that phase separation of the protein PGL-3 is driven by an α-helical region, while the disordered region may stabilize condensates by modulating the driving forces for phase separation.

We next investigated how RNA impacts and alters the phase behaviour of PGL-3. It has been previously shown that the PGL-3 protein cannot phase separate independently at physiological concentration (Saha *et al*, 2016). At this concentration, sufficiently long mRNA can trigger phase separation via binding to six arginine-glycine-glycine (RGG) repeats at the C-terminal disordered region of PGL-3 (Saha *et al*, 2016). We therefore expected that when the C-terminal disordered region was mixed with RNA, it would promote phase separation. Unexpectedly, we found that in contrast to full-length PGL-3, mRNA does not support phase separation of the C-terminal disordered region of PGL-3 (Fig. 2a, b). This suggests that the N-terminal α-helical region is essential for phase separation. However, we found that mRNA does not enhance phase separation of the N-terminal α-helical region of PGL-3 either (Fig. 2a, b).

**Fig. 2.**
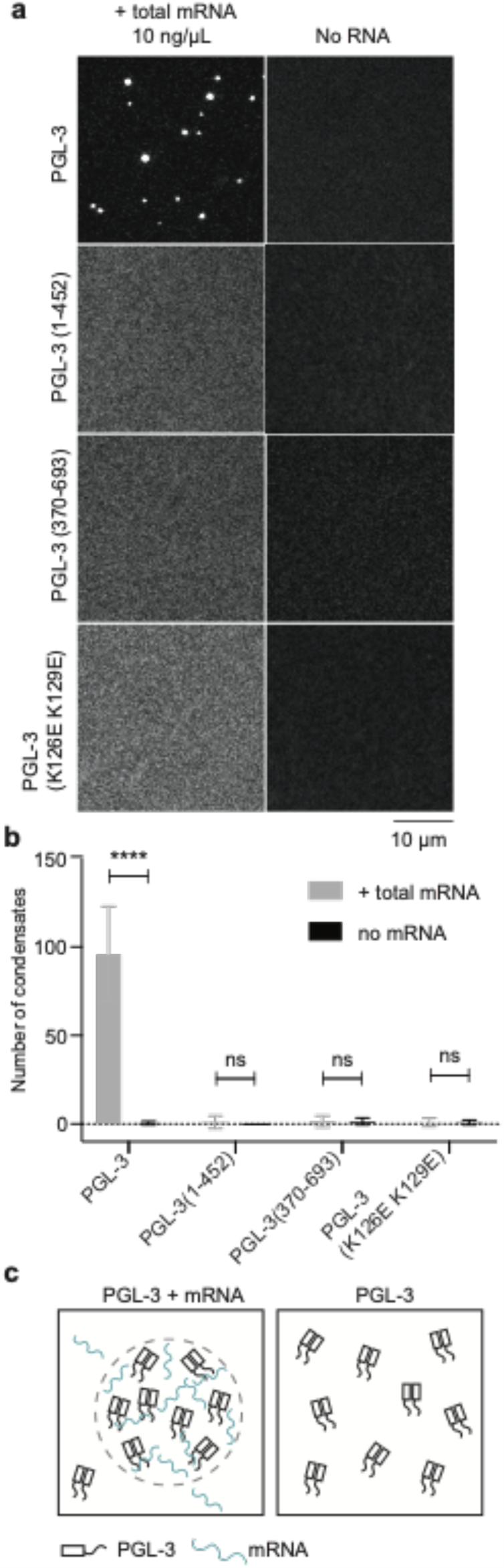
Homotypic interaction among PGL-3 molecules is essential for mRNA-dependent phase separation. **a** Fluorescence micrographs of PGL-3 and mutants (0.5 μM) in presence or absence of total mRNA (10 ng/μL). 5% of the pool of protein was tagged with mEGFP in each case. Buffer: 25 mM HEPES pH 7.5, 150 mM KCl, 1 mM DTT. 1% PEG 35,000 was added to PGL-3 (1-452). Scale bar 10 μm. **b** Plot of the number of condensates with PGL-3 and mutants in presence or absence of total mRNA. For each condition, number of condensates was scored in nine different fluorescence micrographs (116 μm × 117 μm). Error bars represent 1 SD. t-test was performed to compare the number of condensates of each construct with or without the addition of total mRNA (p values: PGL-3: p = 1.04E-6; PGL-3 (1-452): p = 0.172; PGL-3 (370-693): p = 0.940; PGL-3 (K126E K129E): p = 0.828). The significance score was determined by the following criteria: ns: p > 0.05; ****: p ≤ 0.0001. **c** Cartoon showing that at physiological concentrations, mRNA facilitates PGL-3 condensate formation. Association among folded N-termini of PGL-3 molecules is necessary for phase separation.

To investigate the underlying mechanism further, we began by testing if the N-terminal α-helical region of PGL-3 can self-associate. Our analysis using size exclusion chromatography followed by multi-angle light scattering (SEC-MALS) showed that this PGL-3 fragment 1-452 forms a dimer (Supplementary Fig. 2f). Mutation of two residues (K126E K129E) have been shown to interfere with interactions among the N-termini of PGL-3 molecules (Aoki *et al*, 2021). We mutated these two residues within the full-length PGL-3 protein (K126E K129E) (Fig. 1a) and found that this mutant PGL-3 (K126E K129E) protein cannot phase separate even at high protein concentrations up to ~130 µM (Fig. 1b, c). Addition of mRNA does not trigger phase separation of this protein at physiological concentrations either (Fig. 2a, b). Taken together, our data is consistent with a model where association among folded N-termini of PGL-3 molecules is essential for phase separation. At physiological concentrations, mRNA facilitates this process by binding to the disordered C-terminus of PGL-3 (Fig. 2c).

### α-helicity of PGL-3 correlates with dynamics in condensates in vitro

P granules have a complex composition, containing mRNA and over forty proteins (Updike & Strome, 2010). It is unclear how dynamics within the P granule phase depends on interactions among those components. Two constitutive P granule proteins, PGL-1 and GLH-1, have been reported to exchange rapidly between the cytoplasm and liquid-like P granules in *C. elegans* gonads (FRAP half-time (t_1/2_) = 7 – 28 s) (Brangwynne *et al*, 2009). Interestingly, in spite of their vastly reduced compositional complexity, we found that condensates generated in vitro with the PGL-3 protein alone show similar dynamics (t_1/2_ = 7 s) (Supplementary Fig. 2c). To investigate the role of other P granule components on dynamics of PGL-3, we added mRNA and recombinantly purified PGL-1 and GLH-1 to condensates of PGL-3 in vitro at physiologically relevant proportions compared to PGL-3 (Saha *et al*, 2016). We found that PGL-1, GLH-1 and mRNA concentrate within condensates of PGL-3 in vitro (Supplementary Fig. 4d-f), but have a modest effect on the dynamics of PGL-3 (t_1/2_ = 10 s) (Supplementary Fig. 2c). This suggests that the dynamics of full length PGL-3 are intrinsic to the protein, and that the influence of complex composition on dynamics is modest.

In order to address the interactions that drive dynamics within condensates, we began by probing the dynamics of several PGL-3 constructs within phase-separated condensates in vitro. The rate at which molecules of two PGL-3 constructs 1-452 and 1-577, that either lack the disordered C-terminal region entirely or in part, rearrange within condensates are close to full-length PGL-3 (Fig. 1e, Supplementary Fig. 2g, 3a, b). This suggests that interactions involving the disordered C-terminal region of PGL-3 are not essential for the fast dynamics within condensates. Therefore, we addressed the role of the N-terminal α-helical region (1-452) in driving dynamics. In order to avoid engineering mutations that result in significant misfolding of PGL-3 and concomitant loss of its ability to phase separate, we focused our mutational analysis close to the junction of the folded N-terminus and the disordered C-terminus of PGL-3. Surprisingly, we found that a full-length PGL-3 construct (Δ425-452) that lacks only 27 residues phase separates into aggregates (Fig. 3a, c). Sequence analysis of the PGL-3 protein predicts that this region 425-452 spans two α-helices (one complete helix and fraction of a second helix) (Supplementary Fig. 3d). We generated a PGL-3 construct (hereafter called ‘S1’) (Fig. 3a) in which the sequence in the region, 425-452, is shuffled while keeping the overall amino acid composition unchanged. We found that S1 phase separates into condensates that are 20-fold less dynamic than with wild-type PGL-3 (Fig. 3d, Supplementary Fig. 3c). Slow dynamics within condensates is not a result of aggregation of S1 molecules, since condensates of S1 dissolve without detectable trace of aggregates on addition of excess salt (Supplementary Figure 3j). Following assembly via phase separation, the extremely slow dynamics in condensates of S1 does not change significantly over 24 h (Supplementary Fig. 3f). Taken together, our data suggests that interactions involving the N-terminal α-helical region of PGL-3 are required for fast dynamics within condensates.

**Fig. 3.**
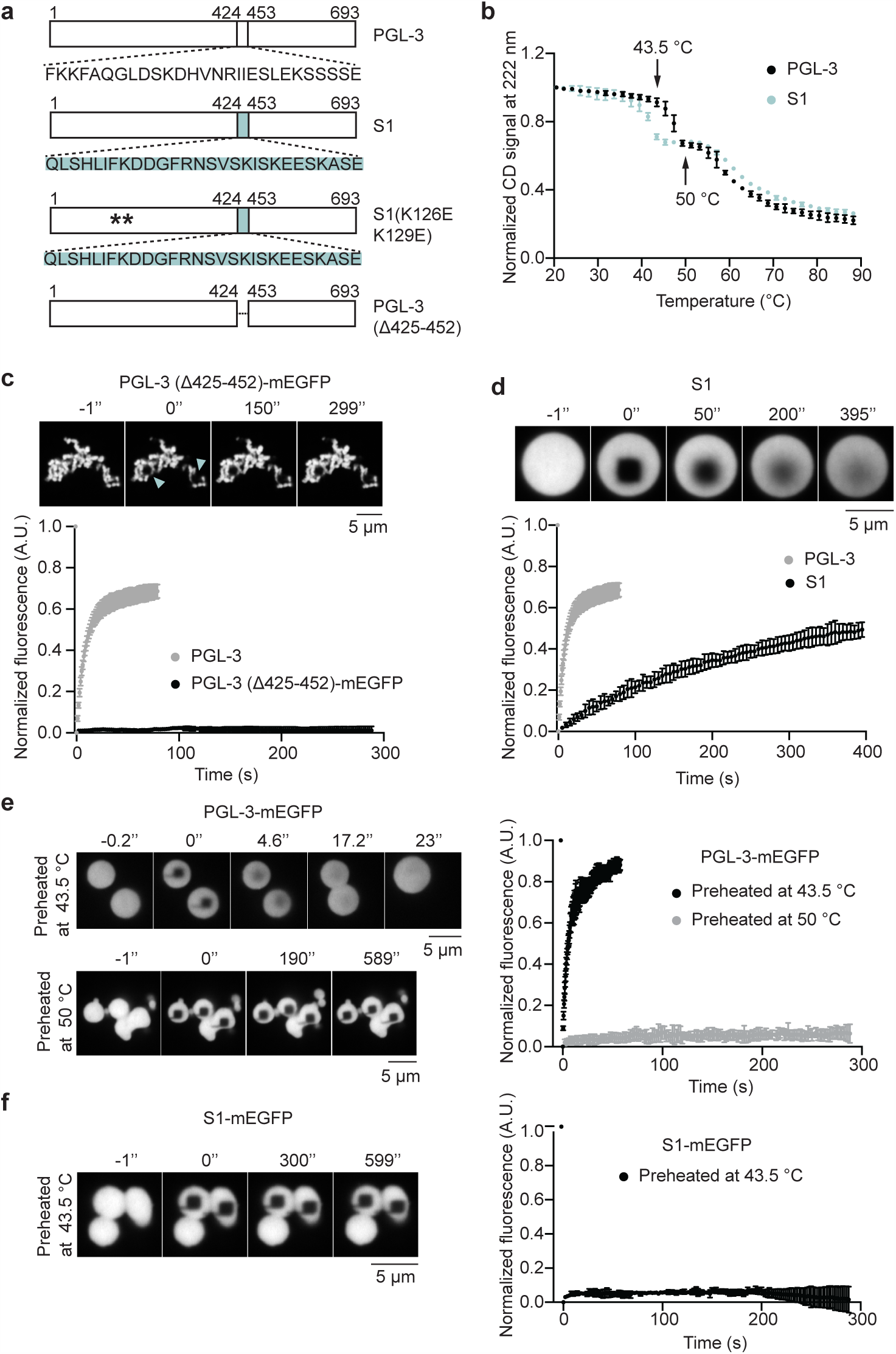
α-helicity of PGL-3 correlates with dynamics in condensates. **a** Cartoon representing PGL-3, S1, S1(K126E K129E) and PGL-3 (Δ425-452). Residues 425-452 of wild-type PGL-3 are shuffled in mutants S1 and S1(K126E K129E). **b** Plot of normalized circular dichroism (CD) spectroscopy signal at 222 nm with PGL-3 (black circle) and S1 (blue circle) vs. temperature, assayed at 5 μM protein concentration in 25 mM HEPES pH 7.5, 1 M KCl, 1 mM DTT. Error bars represent 1 SD among three independent measurements. Arrows highlight 43.5 °C and 50 °C – temperatures where partial unfolding of PGL-3 begins and ends respectively. **c-d** Time-lapse fluorescence micrographs show fluorescence recovery after photobleaching in condensates of **c** PGL-3 (Δ425-452)-mEGFP or **d** S1 (5% of the pool of protein was tagged with mEGFP). Arrowheads show photobleached area in condensates of PGL-3 (Δ425-452)-mEGFP. Scale bars 5 μm. Shown below the time-lapse micrographs are plots of average normalized fluorescence over time **c** in bleached regions (1.4 μm × 1.4 μm) of PGL-3 (Δ425-452)-mEGFP condensates (n = 8) or **d** in bleached regions (1.5 μm × 1.5 μm) of S1 (5% tagged with mEGFP) condensates (n = 18). In both plots, normalized fluorescence over time in bleached regions (1.5 μm × 1.5 μm) of wild-type PGL-3 (5% of the pool of protein tagged with mEGFP) condensates (n = 18) are also shown as reference. Error bars represent 1 SD. **e** Time-lapse fluorescence micrographs show fluorescence recovery after photobleaching at room temperature in condensates of PGL-3-mEGFP (preheated at 43.5 °C or 50 °C). Scale bar 5 μm. Shown next to the micrographs is a plot of average normalized fluorescence over time in bleached regions (1.4 μm × 1.4 μm) within the condensates of PGL-3-mEGFP preheated at 43.5 °C (black circles, n =8) or 50 °C (grey circles, n = 8). Error bars represent 1 SD. **f** Time-lapse fluorescence micrographs show fluorescence recovery after photobleaching at room temperature in condensates of S1-mEGFP (preheated at 43.5 °C). Scale bar 5 μm. Shown next to the micrographs is a plot of average normalized fluorescence over time in bleached regions (1.4 μm × 1.4 μm) within the condensates of S1-mEGFP preheated at 43.5 °C (n =8). Error bars represent 1 SD.

For more insights into the role of the N-terminal α-helical region in regulating dynamics within condensates of PGL-3, we investigated the contribution of the region 425-452 further. CD spectroscopy showed that α-helicity is reduced in the mutant S1 compared to wild-type PGL-3 (Supplementary Fig. 3g). One possibility is that sequence-shuffling of residues 425-452 changes the tertiary structure of PGL-3. Interestingly, we found that the mutant S1 can phase separate at concentrations lower than wild-type PGL-3 (Supplementary Fig. 3e). Further, mutation of two residues (K126E K129E), that interfere with interactions among the N-termini of wild-type PGL-3 molecules (Aoki *et al*, 2021), also inhibit phase separation of the S1 mutant even at high protein concentrations up to ~50 µM (Fig. 3a, Supplementary Fig. 3c, e). This suggests that, regardless of change in tertiary structure, interactions among the N-termini of S1 molecules are important for phase separation. In order to address the consequence of sequence-shuffling of residues 425-452 further, we compared the structural stability of S1 with wild-type PGL-3. We tested this through thermal unfolding which monitor α-helical content between 20 °C and 90 °C, using the CD signal at 222 nm as a measure of α-helical content. We found that wild-type PGL-3 unfolds in two phases as temperature is increased (T_m_ = 47 °C and 61 °C) (Fig. 3b). On the other hand, S1 also unfolds in two phases but sequence-shuffling results in lower thermostability (T_m_ = 41 °C and 61 °C) (Fig. 3b). Taken together, our observations on the phase separation behavior of the S1 mutant raises the possibility that loss of α-helicity of PGL-3 may correlate with slower dynamics in condensates.

In order to address if dynamics in condensates correlate with structural stability of PGL-3, we compared the dynamics of condensates formed via phase separation of folded and unfolded PGL-3 constructs. Heat-induced unfolding of both PGL-3 and S1 is irreversible (Supplementary Fig. 3h, i). These constructs do not recover lost α-helicity once the temperature is lowered to 21 °C following unfolding at higher temperatures. Therefore, we heated mEGFP-tagged PGL-3 and S1 proteins at different temperatures for 10 min followed by cooling to room temperature and triggering phase separation (by diluting salt (KCl) concentration to 150 mM). Wild-type PGL-3 does not lose α-helicity when heated to 43.5 °C (Supplementary Fig. 3h). We observed that PGL-3-mEGFP preheated at 43.5 °C generates condensates that are liquid-like (Fig. 3e). On the other hand, heating S1 to 43.5 °C results in partial unfolding (Supplementary Fig. 3i), and phase separation of this partially unfolded S1-mEGFP protein generates solid-like condensates with no observable FRAP recovery (Fig. 3f). Similarly, heating wild-type PGL-3 to a higher temperature (50 °C) results in partial unfolding (Supplementary Fig. 3h), and phase separation of this partially unfolded PGL-3-mEGFP protein generates solid-like condensates with no observable FRAP recovery (Fig. 3e). Taken together, our data suggests that reduced α-helicity of PGL-3 correlates with slower dynamics in condensates.

### P granule components buffer against reduced dynamics in condensates

We wanted to see if similar changes in dynamics are seen in P granules in *C. elegans* worms. To address this, we injected mEGFP-tagged wild-type PGL-3 or S1 proteins into gonads of adult *C. elegans* and investigated their dynamics. Both PGL-3-mEGFP and S1-mEGFP target to perinuclear P granules (Fig. 4a). Fluorescence recovery after photobleaching experiments showed that S1 exchanges between P granules and cytoplasm at rates similar to wild-type PGL-3 (Fig. 4b, Supplementary Fig 4a). Similarly, when partially unfolded PGL-3-mEGFP (preheated at 50 °C) is injected into gonads of adult *C. elegans*, this protein concentrates in perinuclear P granules (Fig. 4a) and exchanges between P granules and cytoplasm at rates similar to wild-type PGL-3 (Fig. 4b, Supplementary Fig 4a). Experiments with microinjected mutant PGL-3 proteins most likely assay the effect of the P granule phase on high non-physiological dose of partially denatured PGL-3 proteins. To test the effect on a physiologically expressed mutant PGL-3 protein, we used CRISPR/Cas9 technology to generate a transgenic strain that expresses the construct PGL-3 (Δ425-452)-mEGFP from the native *pgl-3* locus. We found that PGL-3 (Δ425-452)-mEGFP concentrates in perinuclear puncta (Fig. 4c) and shows dynamics similar to wild-type PGL-3-mEGFP (Fig. 4e, Supplementary Fig. 4b, Supplementary Movie 1 and 2). Further, we found that fecundity of the strain expressing PGL-3 (Δ425-452)-mEGFP is similar to the strain expressing PGL-3-mEGFP (Fig. 4d). This suggests that the P granule phase can prevent slowdown of dynamics and associated loss-of-function resulting from the expression of the aggregation-prone PGL-3 (Δ425-452)-mEGFP construct. Taken together, our results suggest that the multicomponent P granule phase can prevent slowdown of dynamics associated with reduced α-helicity of PGL-3.

**Fig. 4.**
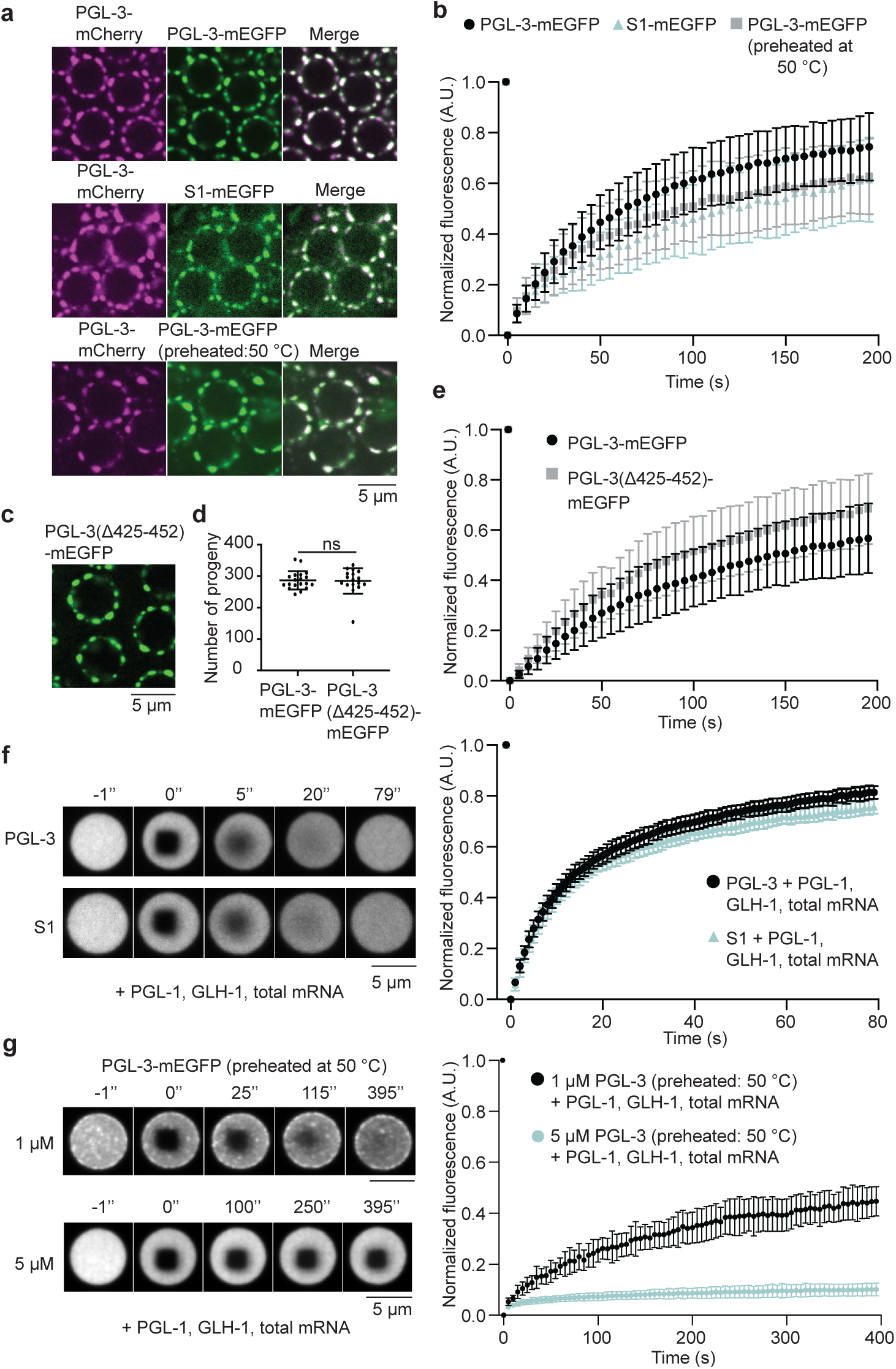
P granule components a promote fast dynamics of PGL-3 constructs with reduced α-helicity. **a** Fluorescence micrographs of gonads (pachytene region) in *C. elegans* expressing PGL-3-mCherry (shown in magenta) injected with recombinantly-purified PGL-3-mEGFP, S1-mEGFP, or PGL-3-mEGFP preheated at 50 °C (shown in green). A single confocal section is shown. Scale bar 5 μm. **b** Plot showing average normalized fluorescence signal over time in bleached peri-nuclear regions (1.5 μm × 1.5 μm) in wild-type *C. elegans* gonads (pachytene region) injected with recombinantly purified PGL-3-mEGFP (black circles, n = 46), S1-mEGFP (blue triangles, n = 36), or PGL-3-mEGFP preheated at 50 °C (grey circles, n = 37). Error bars represent 1 SD. For each condition, data from 2 different animals injected with a recombinantly purified protein were analyzed. Statistical analysis of the estimated half times of fluorescence recovery is shown in Supplementary Fig. 4a. **c** Fluorescence micrograph of gonad (pachytene region) in *C. elegans* expressing PGL-3 (Δ425-452)-mEGFP (shown in green). A single confocal section is shown. Scale bar 5 μm. **d** Plot showing total number of progeny of worms expressing PGL-3-mEGFP (n=19) or PGL-3 (Δ425-452)-mEGFP (n=17) at 20 °C. Each black circle represents the number of progenies from a single worm. The mean and 1 SD are also indicated. For statistical analysis, T test was performed (p = 0.858). “ns” indicates no significant difference (p > 0.05) between the number of progenies of the two genotype. **e** Plot showing average normalized fluorescence signal over time in bleached peri-nuclear regions in *C. elegans* gonads (pachytene region) expressing either PGL-3-mEGFP (black circles, n = 43) or PGL-3 (Δ425-452)-mEGFP (grey squares, n = 32). Error bars represent 1 SD. For each condition, data from 6 different animals were analyzed. Statistical analysis of the estimated half times of fluorescence recovery is shown in Supplementary Fig. 4b. **f** Time-lapse fluorescence micrographs show fluorescence recovery after photobleaching over time in condensates of PGL-3 (5 μM) or S1 (5 μM) (5% of the pool of protein was tagged with mEGFP) in presence of PGL-1 (6.7 μM), GLH-1 (1 μM), and total mRNA (14 ng/μL). Scale bar 5 μm. Shown next to the micrographs is a plot of average normalized fluorescence over time in bleached regions (1.5 μm × 1.5 μm) within the condensates of PGL-3 (black circles, n = 20) or S1 (blue triangles, n = 21) in presence of PGL-1, GLH-1 and total mRNA. Error bars represent 1 SD. Statistical analysis of the estimated half times of fluorescence recovery is shown in Supplementary Fig. 4c. **g** Time-lapse fluorescence micrographs show fluorescence recovery after photobleaching over time in condensates of PGL-3-mEGFP preheated at 50 °C in presence of PGL-1 (6.7 μM), GLH-1 (1 μM), and total mRNA (14 ng/μL). Concentration of PGL-3-mEGFP used: 1 μM or 5 μM. Scale bar 5 μm. Shown next to the micrographs is a plot of average normalized fluorescence over time in bleached regions (1.5 μm × 1.5 μm) within the condensates of PGL-3-mEGFP preheated at 50 °C. Black circles, 1 μM PGL-3-mEGFP (n = 14) and blue circles, 5 μM PGL-3-mEGFP (n = 15). Error bars represent 1 SD.

To reconstitute the multicomponent nature of the P granule, we purified and added back other P granule proteins to the PGL-3 phase formed in vitro. Interestingly, addition of merely a few P granule components (mRNA and recombinantly purified PGL-1 and GLH-1) to condensates of S1 in vitro enhances the dynamics of S1 almost 20-fold to match the dynamics observed in condensates of wild-type PGL-3 (Fig. 4f, Supplementary Fig. 4c-e, 5f). Further, we found that dynamics within condensates of S1 can be significantly enhanced simply by adding wild-type PGL-3 protein (Supplementary Fig. 5a). For better insight into how the PGL-3 protein supports fast dynamics of S1, we added three different PGL-3 constructs (the N-terminal α-helical fragment 1-452, the C-terminal fragment 370-693, and the phase separation-incompetent mutant PGL-3 (K126E K129E)) at different concentrations to condensates of S1. We found that all three PGL-3 constructs concentrate within condensates of S1 to different degrees (Supplementary Figure 5c) and enhance the dynamics of S1 (Supplementary Fig. 5a). Further, addition of the phase separation-incompetent mutant S1 (K126E K129E) to condensates of S1 also enhanced the dynamics of S1 within condensates (Supplementary Fig. 5a and 5c). Taken together, our data suggests that enhanced S1 dynamics in condensates can be supported by a wide variety of interactions involving other P granule components (PGL-1, GLH-1, mRNA) or different parts of the PGL-3 protein.

We next wondered if addition of other P granule components is able to support or rescue a dynamic state of partially unfolded PGL-3 within condensates. In contrast to S1, which generates condensates that are liquid-like, partially unfolded PGL-3-mEGFP (preheated at 50 °C) phase separates into non-dynamic solid-like condensates in vitro (Fig. 3e). We tested the ability of different components to rescue dynamics of partially unfolded PGL-3-mEGFP. When P granule components (mRNA and recombinantly purified PGL-1 and GLH-1) are added to partially unfolded PGL-3-mEGFP at concentrations similar to those used with S1 (5 μM), the PGL-3 protein phase separates predominantly into non-dynamic condensates (Fig. 4g, Supplementary Fig. 4f, g). However, when a lower concentration of the partially unfolded PGL-3 protein is used (1 μM), addition of these P granule components supports its phase separation into liquid-like condensates (Fig. 4g). These results revealed a concentration-dependence to the extent with which unfolded PGL-3 impacts condensate dynamics.

We wondered if, similar to partially unfolded PGL-3-mEGFP, the dynamics of S1 also depends on its concentration. In the assays in which S1 is mixed with different additional components, 5% of the total pool of S1 is tagged with mEGFP. This allowed us to measure the dynamics of S1 within condensates using FRAP. Following photobleaching, fluorescence recovered in bleached region within condensates in two phases – a fast phase driven predominantly by internal rearrangement of molecules within the condensate, and a slower phase driven by the exchange with the pool of unbleached molecules outside the condensate (Supplementary Fig. 6a-f). We focused on the fast phase of recovery and used single-exponential fits to estimate half-time of internal rearrangement of S1 molecules within condensates of different composition. On addition of other P granule components (PGL-1, GLH-1, mRNA) to different concentrations of S1, we found that the half-time of FRAP recovery of S1 within condensates remains relatively unchanged in spite of almost 5-fold change in mEGFP-fluorescence signal per unit area within the condensates (Fig. 5a). Assuming that mEGFP-fluorescence reports on the concentration of S1, this observation suggests that P granule components can buffer the dynamics of S1 close to rates observed in condensates of wild-type PGL-3 alone (Fig. 5a, d). However, the dynamics of S1 slows down above a threshold concentration (9 μM in Fig. 5a), suggesting that P granule components can buffer the dynamics of S1 only within a range of concentrations in condensates. We wondered if the observed dynamics buffering effect within condensates of S1 is limited to the specific set of additives PGL-1, GLH-1 and mRNA. We assayed this by adding to condensates of S1 the construct PGL-3 (K126E K129E). The mutations K126E K129E are expected to interfere with interactions among N-termini of S1 and PGL-3 (K126E K129E) molecules (Aoki *et al*, 2021). We found that addition of this phase separation-incompetent mutant PGL-3 (K126E K129E) can also buffer dynamics within a 2-fold range of S1 concentration within condensates (Fig. 5b, Supplementary Fig. 5g). However, the buffering is different compared to when PGL-1, GLH-1 and mRNA are added. Specifically, the range of concentrations of S1 in condensates where dynamics is buffered and the associated half-time of FRAP recovery depend on the additives. We used condensates of similar size for our analysis (average ± 1 SD of diameter of condensates are 6.4 ± 1.7 μm (Fig. 5a) and 5.9 ± 0.4 μm (Fig. 5b)). Therefore, dynamics buffering here is likely to represent similar diffusion rates of S1 within condensates.

**Fig. 5.**
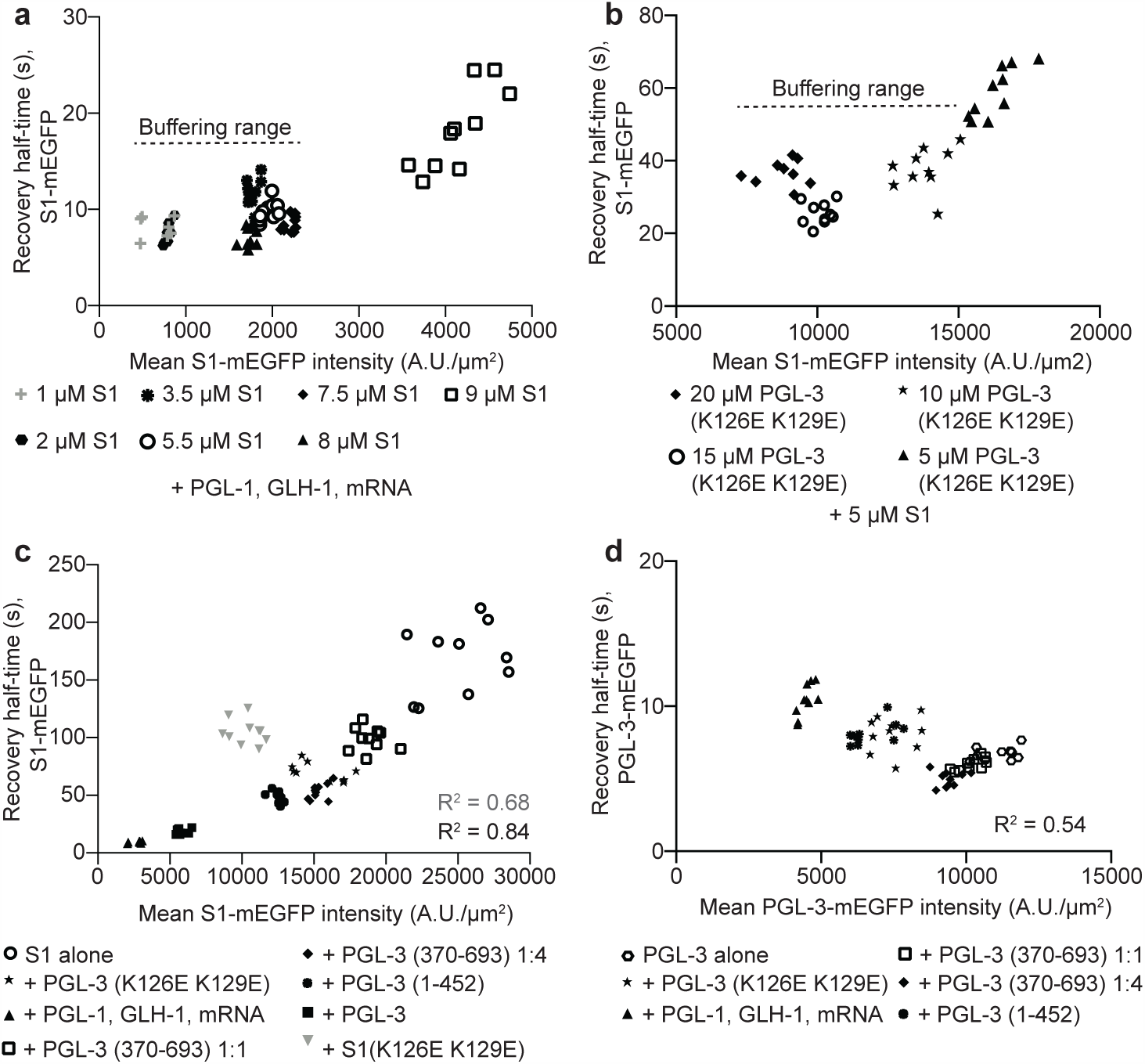
Dynamics of S1 in condensates of different composition. Analysis of fluorescence recovery after photobleaching within 1.5 μm × 1.5 μm regions in condensates of different composition. Half-time of fluorescence recovery after photobleaching (t_1/2_) in each case was estimated by exponential curve (see methods section for details). **a** Plot of t_1/2_ vs. mean S1-mEGFP fluorescence intensity per unit area (before photobleaching) within condensates. Each point on the plots represents data from a single photobleached region within a condensate of defined composition which includes 5 μM PGL-1, 0.75 μM GLH-1, 11 ng/ μM mRNA and different S1 concentrations indicated as follows. Grey plus: 1 μM S1, solid hexagon: 2 μM S1, asterisk 3.5 μM S1, open circle: 5.5 μM S1, solid diamond: 7.5 μM S1, solid triangle: 8 μM S1, open square: 9 μM S1. Ten points are plotted for each composition (seven for 1 μM S1) and buffering range indicates a range of S1 concentrations where t_1/2_ does not differ significantly between different conditions. **b** Plot of t1/2 vs. mean S1-mEGFP fluorescence intensity per unit area (before photobleaching) within condensates. Each point on the plots represents data from a single photobleached region within a condensate of defined composition which includes 5 μM S1 and different concentrations of PGL-3 (K126E K129E) indicated as follows. Solid diamond: 20 μM PGL-3 (K126E K129E), open circle: 15 μM PGL-3 (K126E K129E), asterisk 10 μM PGL-3 (K126E K129E), solid triangle: 5 μM PGL-3 (K126E K129E), Ten points are plotted for each composition (nine for 20 μM PGL-3 (K126E K129E)) and buffering range indicates a range of S1 concentrations where t1/2 does not differ significantly between different conditions. **c** Plot of t_1/2_ vs. mean S1-mEGFP fluorescence intensity per unit area (before photobleaching) within condensates. Each point on the plots represents data from a single photobleached region within a condensate of defined composition as follows. Open circle: S1 alone (10 μM, 5% mEGFP-tagged), solid diamond: S1 (5 μM, 5% mEGFP-tagged) + PGL-3 (370-693) (20 μM), solid asterisk: S1 (5 μM, 5% mEGFP-tagged) + PGL-3 (K126E K129E) (5 μM), solid circle: S1 (5 μM, 5% mEGFP-tagged) + PGL-3 (1-452) (5 μM), solid triangle: S1 (5 μM, 5% mEGFP-tagged) + PGL-1 (6.7 μM) + GLH-1 (1 μM) + total mRNA (14 ng/μL), solid square: S1 (5 μM, 5% mEGFP-tagged) + PGL-3 (5 μM), open square: S1 (5 μM, 5% mEGFP-tagged) + PGL-3 (370-693) (5 μM), grey inverse triangle: S1 (5 μM, 5% mEGFP-tagged) + S1(K126E K129E) (5 μM). Ten points are plotted for each composition. All or part of the data were fit using linear regression (R^2^ = 0.68, fit of all data; R^2^ = 0.84, fit of data on all compositions excluding S1+ S1(K126E K129E)). Data points for S1+ S1(K126E K129E) are highlighted since S1(K126E K129E) is the only construct added to S1 which contains the shuffled 425-452 region. **d** Plot of t_1/2_ vs. mean PGL-3-mEGFP fluorescence intensity per unit area (before photobleaching) within condensates. Each point on the plots represents data from a single photobleached region (1.5 μm × 1.5 μm) within a condensate of defined composition as follows. Open hexagon: PGL-3 alone (10 μM, 5% mEGFP-tagged), open square: PGL-3 (5 μM, 5% mEGFP-tagged) + PGL-3 (370-693) (5 μM), solid asterisk: PGL-3 (5 μM, 5% mEGFP-tagged) + PGL-3 (K126E K129E) (5 μM), solid diamond: PGL-3 (5 μM, 5% mEGFP-tagged) + PGL-3 (370-693) (20 μM), solid triangle: PGL-3 (5 μM, 5% mEGFP-tagged) + PGL-1 (6.7 μM) + GLH-1 (1 μM) + total mRNA (14 ng/μL), solid circle: PGL-3 (5 μM, 5% mEGFP-tagged) + PGL-3 (1-452) (5 μM). Ten points are plotted for each composition and R^2^ value for a linear regression fit among all compositions is indicated.

For more insights into how addition of different components rescues slowdown of S1 dynamics within condensates, we added wild-type PGL-3, or different PGL-3 or S1 constructs to condensates of S1. These additives titrated the concentration of S1 within condensates (tracked by S1-mEGFP fluorescence signal per unit area) over a range spanning concentration observed with condensates of S1 alone and those in presence of other P granule components (PGL-1, GLH-1, mRNA). Here, we found that the half-time of FRAP recovery of S1 correlates with mEGFP-fluorescence signal per unit area within the condensates (Fig. 5c). This suggests that, regardless of the condensate composition, fast dynamics generally correlate with smaller concentrations of S1. In control assays, we found that the dynamics of the wild-type PGL-3 protein does not correlate significantly with either concentration of PGL-3 or the composition of the condensate (Fig. 5d, Supplementary Fig. 5b, d). While fast dynamics in condensates generally correlate with smaller concentrations of S1 molecules (Fig. 5c), the influence of relative concentrations of macromolecules within condensates on dynamics buffering remains unclear. To address this, we estimated the ratio of intra-condensate concentrations of S1 and PGL-3 (K126E K129E) both within and outside the ‘buffering range’ (see Fig. 5b). We found that dynamics of S1 molecules is buffered when the intra-condensate concentration of the additive PGL-3 (K126E K129E) is roughly equal to or exceeds the concentration of S1 within condensates (Supplementary Fig. 5h).

## Discussion

To maintain homeostasis, cellular liquid-like compartments must be functionally robust against small changes in composition or denaturation of one or more of its components. The mechanisms that provide such robustness remain unclear. A recent study proposed that liquid-liquid phase separation may reduce variation in concentration of components within condensates (Klosin *et al*, 2020). It remains unclear if LLPS can similarly buffer diffusion rate of components within condensates. Recent studies with simple two-component LLPS systems showed that the behavior of one component can influence the dynamics of another within condensates (Boeynaems *et al*, 2019; Rhine *et al*, 2020; Zhang *et al*, 2015; Alshareedah *et al*, 2021). While components are expected to influence the dynamics of each other within multi-component condensates, it is unclear *a priori* how dynamics of biomolecular condensates with complex composition adapt to misfolding/denaturation of one or more component proteins. Here, we show that the liquid-like P granule phase can rescue slow-down of diffusion rate associated with denaturation of a component protein. Such dynamics-buffering may contribute to functional robustness of cellular liquid-like compartments.

In order to understand how composition supports dynamics-buffering in biomolecular condensates, we began by addressing why denaturation of the PGL-3 protein leads to slower dynamics in condensates. One possibility is that the conformational changes of PGL-3 associated with reduced α-helicity facilitate stronger inter-molecular interactions. For instance, loss of α-helicity could increase the valence of “stickers” leading to additional physical crosslinks among PGL-3 molecules (Choi *et al*, 2020). Stronger inter-molecular interactions could inhibit rapid rearrangement of interacting partners within condensates, resulting in slower dynamics. Alternatively, interactions involving unfolded or partially unfolded PGL-3 molecules could generate inter-polypeptide chain entanglements within condensates resulting in slower dynamics. With diffusion of PGL-3 molecules within condensates, more entanglements could develop over time resulting in further decrease in dynamics. Few observations suggest that slower dynamics within PGL-3 condensates result predominantly from stronger inter-molecular interactions in contrast to inter-chain entanglements. First, dynamics of the mutant S1 protein within condensates does not change significantly over 24 h. Second, the mutations K126E K129E that interfere with interactions among PGL-3 or S1 molecules (Aoki *et al*, 2021) help enhance dynamics of S1 molecules within condensates. For instance, when added to condensates of S1, both S1 (K126E K129E) and PGL-3 (K126E K129E) help enhance the dynamics of S1 within condensates. Addition of S1 (K126E K129E) reduces concentration of S1 within condensates more than addition of PGL-3 (K126E K129E). However, dynamics of S1 within condensates is faster in spite of higher concentration with PGL-3 (K126E K129E). Taken together, these observations support the model that slow dynamics of S1 within condensates depend on interactions among S1 molecules and suggest that interaction among S1 molecules is stronger than that between wild-type PGL-3 molecules. Stronger interaction among S1 molecules is further supported by the observation that S1 molecules can phase separate at lower concentrations compared to wild-type PGL-3 molecules (Supplementary Fig. 3e).

Our data is consistent with the model that other regions of S1 molecules cooperate with residues 425-452 (shuffled) to generate stronger inter-molecular interactions. For instance, addition of the mutant S1 (K126E K129E) enhances dynamics of S1 within condensates in contrast to maintaining the slower dynamics observed within condensates of S1 alone. This suggests that the interactions disrupted by the mutations K126E and K129E also contribute to slow S1 dynamics. One possibility is that interactions involving the residues K126 and K129 favor S1 conformations that enhance 425-452 (shuffled)-dependent interactions. Indeed, the mutations K126E K129E have been reported to interfere with interactions among N-termini of PGL-3 molecules (Aoki *et al*, 2021). While two self-association domains within the α-helical N-terminus of PGL-3 have been mapped (Aoki *et al*, 2021, 2016), structural insights into those associations are limited. However, PGL-3 shares significant sequence similarity with another protein PGL-1. Crystal structures are available for fragments of the PGL-1 protein that show the two self-association domains at the N-terminus are predominantly α-helical and globular in nature (Aoki *et al*, 2016, 2021). Therefore, one possibility is that shuffling the sequence 425-452 of PGL-3 or heat-induced unfolding of PGL-3 exposes hydrophobic residues that become available to participate in inter-molecular interactions. This effect may be enhanced within phase separated condensates if the local solvent environment favors more extended conformations of the PGL-3 protein, allowing more inter-molecular interactions compared to within the surrounding dilute phase. Indeed, recent studies reported that a 129-residue long intrinsically disordered fragment of the human Tau protein or the prion-like low complexity domain of hnRNPA1 preferentially sample more extended conformations within phase-separated condensates compared to the surrounding dilute phase (Majumdar *et al*, 2019; Bremer *et al*, 2021). Taken together, our data suggests that strength of inter-molecular interactions regulate dynamics within condensates of PGL-3.

It is generally thought that chaperone proteins or RNA helicases counteract slow-down of dynamics that may result from denaturation of components or inter-polypeptide/RNA chain entanglements in biomolecular condensates. For instance, the ATPase-dependent activity of chaperones is required to dissolve aberrant stress granules that result from aggregation of misfolding-prone proteins (Ganassi *et al*, 2016; Mateju *et al*, 2017). In contrast, we found that weak interactions among two or more components can buffer against large change of dynamics within the P granule phase. This effect is distinct from active processes that depend on ATP hydrolysis. Specifically, we show that presence of just three P granule components (PGL-1, GLH-1 and mRNA) is sufficient to support a dynamic state of partially unfolded PGL-3-mEGFP within phase-separated condensates. In absence of the P granule components, partially unfolded PGL-3-mEGFP phase separates into non-dynamic condensates. We found that the ability of P granule components to support a dynamic state of partially unfolded PGL-3-mEGFP depends on the relative concentrations and hence stoichiometries of PGL-3 and the P granule components. This suggests that P granule components support dynamics by participating in inter-molecular interactions with PGL-3-mEGFP molecules. Similarly, different PGL-3 constructs – PGL-3 (K126E K129E), PGL-3 (1-452) and PGL-3 (370-693) enhanced dynamics of S1 molecules within condensates to different degrees. This may correlate with the varying ability of these constructs to interact with S1 molecules. We considered the specificity of interactions that support fast dynamics within phase separated condensates. Our data showing that different additives (P granule components, or wild-type PGL-3, or PGL-3 constructs that differ in terms of their ability to fold or self-associate) can all rescue slowdown of dynamics of the mutant S1 protein within condensates suggest that fast dynamics can be supported by competing interactions among two or more components (Fig. 6). Future investigations will reveal the individual contribution of different P granule proteins to dynamics buffering. Similar to our data with P granules, interactions among components could be a potential mechanism of storage of misfolding-prone proteins in non-aggregated state within the liquid-like nucleolus under stress in vivo (Frottin *et al*, 2019). We speculate that these interactions that support fast dynamics are weak in nature and allow fast dissociation rates. Taken together, our study supports the possibility that weak interactions among two or more components prevent aggregation or large change of dynamics within cellular liquid-like compartments.

**Fig. 6.**
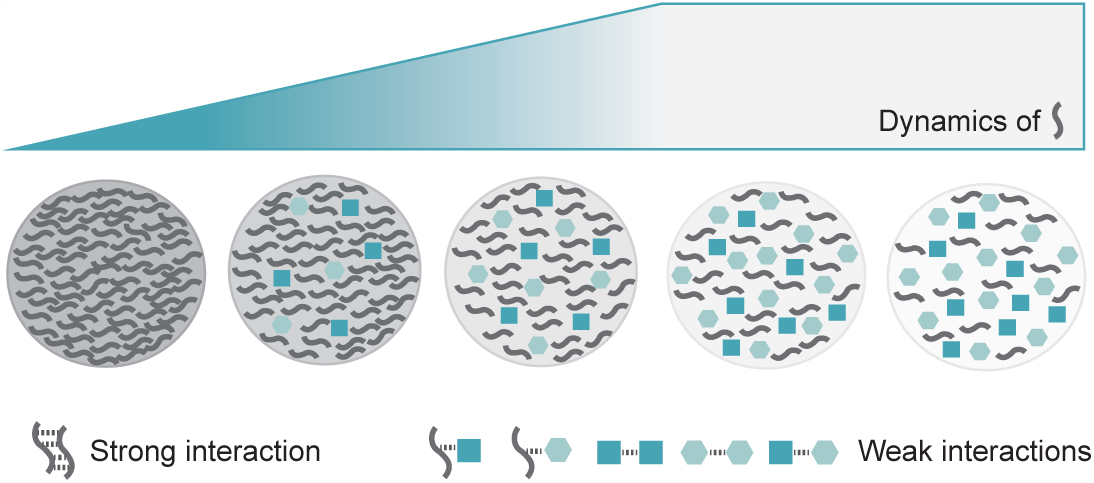
Weak interactions among two or more components promote fast dynamics in condensates. Cartoon shows how dynamics in condensates change with composition. Each condensate contains one or more of three different macromolecules (depicted as thread, square and hexagon) at different concentrations. Homotypic interactions among threads are strong. Other homotypic and heterotypic interactions are weak. Dynamics generally correlate inversely with concentration of strong interactors in condensates. However, within a range of low concentrations of strong interactors, heterotypic interactions can buffer, at least in part, against significant change of dynamics.

An interesting aspect of the dynamics buffering effect is that the dynamics of S1 remain relatively unchanged regardless of the change of S1 concentration within condensates. The buffering capacity depends on the additives i.e. the range of intra-condensate S1 concentrations where dynamics is buffered varies with additives. Once the buffering capacity is exhausted i.e. beyond a threshold intra-condensate S1 concentration, dynamics of S1 molecules slow down at higher S1 concentrations. More work is required to understand the underlying mechanism. The “buffered” dynamics of S1 molecules within condensates also vary with additives. With the additive PGL-3 (K126E K129E), we found that dynamics of S1 molecules is buffered when the intra-condensate concentration of the additive is roughly equal to or exceeds the concentration of S1 within condensates (Supplementary Fig. 5h). This is consistent with the notion that weak heterotypic interactions with PGL-3 (K126E K129E) molecules mediate dynamics buffering in condensates of S1. Interestingly, we found that buffered dynamics of S1 molecules is faster in presence of P granule components (PGL-1, GLH-1 and mRNA) compared to that with the mutant PGL-3 (K126E K129E) (compare Fig. 5a and 5b). This difference in buffered dynamics is unlikely simply due to difference in relative concentrations of S1 and additives within condensates. We found that, with the additive PGL-3 (K126E K129E), the dynamics of S1 within condensates remains buffered despite almost 3-fold variation in the relative concentration of S1 and additive (Supplementary Fig. 5h). Future work will reveal if the buffered dynamics depend on the number of interacting species within condensates (e.g. four when PGL-1, GLH-1 and mRNA is added to condensates of S1; two when PGL-3 (K126E K129E) is added to S1 condensates) in addition to the strength/nature of inter-molecular interactions.

Many cellular non-membrane-bound compartments have been characterized to contain two or more phases with distinct biophysical properties (Banani *et al*, 2017; Feric *et al*, 2016; Jain *et al*, 2016; Lyon *et al*, 2021; Wang *et al*, 2014). Further, within the same liquid-like phase, diffusion rates of two different components may differ from each other (Boeynaems *et al*, 2019; Mathieu *et al*, 2020; Zhang *et al*, 2015). In P granules of *C. elegans* early embryos, a gel-like phase of MEG proteins is known to associate with the liquid-like phase of PGL proteins (Wang *et al*, 2014; Smith *et al*, 2016; Putnam *et al*, 2019). The gel-like phase of MEG proteins is absent from P granules of adult *C. elegans* gonads. Two recent studies suggest that association of the gel-like MEG phase changes the surface tension of the liquid-like PGL phase without changing its viscosity or the diffusion rates of PGL proteins (Folkmann *et al*, 2021; Putnam *et al*, 2019). Taken together, one possibility is that the dynamics-buffering mechanism that we identified here adds stability to the steady-state dynamics of liquid-like phases.

An important part of our study is to show that interactions among the α-helical N-terminal domain of PGL-3 molecules can drive phase separation. Although the unstructured C-terminus is dispensable for phase separation of PGL-3, it fine-tunes the process. First, full-length PGL-3 can phase separate at concentrations lower than the α-helical N-terminus 1-452 alone (Fig. 1c), suggesting that interactions involving the C-terminus enhance the propensity of phase separation. Second, heterotypic interactions among the unstructured C-terminus of PGL-3 molecules and mRNA facilitate phase separation of PGL-3 (Saha *et al*, 2016). Our data is consistent with other studies which found that interactions involving both folded domains and intrinsically disordered regions of proteins drive liquid-liquid phase separation to assemble many cellular non-membrane-bound compartments (Guillén-Boixet *et al*, 2020; Mittag & Parker, 2018; Yang *et al*, 2020).

The concept of robustness is widely understood to be essential for the ability of genetic circuits to function under a wide variety of perturbations (Félix & Barkoulas, 2015). We propose that one route to achieving robustness is through condensates that contribute to spatial and temporal regulation over biochemical reactions. The compositional heterogeneity-dependent dynamics-buffering mechanism that we identified here could add robustness to biochemical reactions within condensates against the deleterious consequences of stresses, short-term effects of chemical modifications, intrinsic mutational changes, and the problems that arise from accumulating high local concentrations of sticky molecules. The ability to buffer against these changes to dynamics not only provides robustness, but these networks of interactions could also provide a source of variation that enable responsiveness when conditions change.

## Materials and Methods

### Protein expression and purification

All PGL-3, GLH-1 and PGL-1 constructs were purified recombinantly from insect cells using a baculovirus infection system. Insect cells were harvested ~72 hours after viral infection and lysed. Constructs were purified from lysates using a combination of Ni-NTA affinity, anion exchange and/or size exclusion chromatography as described below.

*PGL-3, PGL-1, PGL-3 (370-693) and PGL-3 (K126E K129E):* The protocol for purification of PGL-3-6xHis-mEGFP has been described earlier (Saha et al., 2016). PGL-3-6xHis-RFP, PGL-1-6xHis-mEGFP, PGL-3 (370-693)-6xHis-mEGFP and PGL-3 (K126E K129E)-6xHis-mEGFP were purified using the same protocol.

*PGL-3 (1-452):* Insect cells expressing PGL-3 (1-452)-6xHis-mEGFP were resuspended in lysis buffer (25 mM HEPES pH 7.5, 300 mM KCl, 0.3 M L-Arginine, 10 mM Imidazole, 1 mM DTT added with protease inhibitor cocktail Complete EDTA-free (Roche) and lysed using sonicator. The lysates were centrifuged at 21,000 × g in a JS-13.1 rotor (Beckman-Coulter), and Ni-NTA Agarose resin (Macherey-Nagel) was mixed with the supernatant to capture PGL-3 protein molecules. The resin was then washed with 25 mM HEPES pH 7.5, 0.3 M KCl, 20 mM Imidazole. Finally, the PGL-3 molecules were eluted into 25 mM HEPES pH 7.5, 300 mM KCl, 250 mM Imidazole.

In preparation for anion-exchange chromatography, the eluate from Ni-NTA column was diluted 1:5 in 25 mM Tris pH 8.0, 1 mM DTT and 5 ml of Q Sepharose resin (GE Healthcare) was mixed to capture PGL-3 protein molecules. After incubation for ~1 h, the resin was washed with 25 mM Tris pH 8.0, 1 mM DTT buffers containing 350 mM, 500 mM and 1 M KCl. The eluates in 25 mM Tris pH 8.0, 1 mM DTT buffers containing 500 mM KCl and 1 M KCl were pooled together and further purified using size exclusion chromatography with a TSKgel 5000PW column (Tosoh Bioscience) in 25 mM HEPES pH 7.5, 1 M KCl, 1 mM DTT.

*PGL-3 (1-577):* Insect cells expressing PGL-3 (1-577)-6xHis-mEGFP were resuspended in lysis buffer (25 mM HEPES pH 7.25, 300 mM KCl, 10 mM Imidazole, 1 mM DTT added with protease Inhibitor cocktail set III (Calbiochem) and lysed via passage through Avestin Emulsiflex C5. The lysates were centrifuged at 20000 rpm in a Ti45 rotor (Beckman-Coulter), and Ni-NTA Agarose resin (Qiagen) was mixed with the supernatant to capture PGL-3 protein molecules. The resin was then washed with 25 mM HEPES pH 7.25, 300 mM KCl, 20 mM Imidazole, 1 mM DTT. Finally, the PGL-3 molecules were eluted into 25 mM HEPES pH 7.25, 300 mM KCl, 250 mM Imidazole.

In preparation for anion-exchange chromatography, the eluate from Ni-NTA column was diluted 1:6 in 25 mM Tris pH 8.0, 1 mM DTT and 5-10 ml of Q Sepharose resin (GE Healthcare) was mixed to capture PGL-3 protein molecules. After incubation for ~1 h, the resin was washed with 25 mM Tris pH 8.0, 1 mM DTT buffers containing 100 mM, 200 mM, 350 mM, 500 mM and 1 M KCl. The PGL-3 molecules eluted into the 25 mM Tris pH 8.0, 1 mM DTT buffers containing 500 mM KCl and 1 M KCl.

The eluates were further purified using size exclusion chromatography with HiLoad 16/60 Superdex 200 column (GE Healthcare) in 25 mM HEPES pH 7.5, 0.3 M KCl, 1 mM DTT. The fractions containing pure PGL-3 (1-577)-6xHis-mEGFP from both runs were pooled together.

*S1 and S1(K126E K129E):* Insect cells expressing PGL-3 (S1)-6xHis-mEGFP or PGL-3 (S1(K126E K129E))-6xHis-mEGFP were resuspended in lysis buffer (25 mM HEPES pH 7.5, 300 mM KCl, 300 mM L-Arginine, 10 mM Imidazole, 1 mM DTT added with protease inhibitor cocktail Complete EDTA-free (Roche) and lysed using sonicator. The lysates were centrifuged at 21,000 × g in a JS-13.1 rotor (Beckman-Coulter), and Ni-NTA Agarose resin (Macherey-Nagel) was mixed with the supernatant to capture PGL-3 protein molecules. The resin was then washed with 25 mM HEPES pH 7.5, 1 M KCl, 20 mM Imidazole. Finally, the PGL-3 molecules were eluted into 25 mM HEPES pH 7.5, 1 M KCl, 250 mM Imidazole. Next, 500 units of Benzonase nuclease and 1 mM each of DTT and MgCl_2_ were added to the eluate and dialyzed overnight into 25 mM HEPES pH 7.5, 1 M KCl, 1 mM DTT, 1 mM MgCl_2_.

In preparation for anion-exchange chromatography, the dialysate was diluted 1:13 in 25 mM Tris pH 8.0, 1 mM DTT and 5-10 ml of Q Sepharose resin (GE Healthcare) was mixed to capture PGL-3 protein molecules. After incubation for ~1 h, the resin was washed with 25 mM Tris pH 8.0, 350 mM KCl, 1 mM DTT and finally the PGL-3 molecules were eluted into 25 mM Tris pH 8.0, 1 M KCl, 1 mM DTT.

The eluates were further purified using size exclusion chromatography with a TSKgel 5000PW column (Tosoh Bioscience) in 25 mM HEPES pH 7.5, 1 M KCl, 1 mM DTT.

*PGL-3 (Δ425-452):* Insect cells expressing PGL-3 (Δ425-452)-6xHis-mEGFP were resuspended in lysis buffer (25 mM HEPES pH 7.25, 300 mM KCl, 300 mM L-Arginine, 10 mM Imidazole, 1 mM DTT, 10 mM MgCl_2_ added with protease Inhibitor cocktail set III (Calbiochem) and Benzonase nuclease) and lysed via passage through Avestin Emulsiflex C5. The lysates were centrifuged at 20000 rpm in a Ti45 rotor (Beckman-Coulter), and Ni-NTA Agarose resin (Qiagen) was mixed with the supernatant to capture PGL-3 protein molecules. The resin was then washed with 25 mM HEPES pH 7.25, 1 M KCl, 40 mM Imidazole, 1 mM DTT. Finally, the PGL-3 molecules were eluted into 25 mM HEPES pH 7.25, 1 M KCl, 250 mM Imidazole, 1 mM DTT. Next, ~250 units of Benzonase nuclease and 10 mM MgCl_2_ were added to the eluate and dialyzed overnight into 25 mM HEPES pH 7.25, 1 M KCl, 1 mM DTT, 10 mM MgCl_2_.

In preparation for anion-exchange chromatography, the dialysate was diluted 1:12 in 25 mM Tris pH 8.0, 1 mM DTT and 5-10 ml of Q Sepharose resin (GE Healthcare) was mixed to capture PGL-3 protein molecules. After incubation for ~1 h, the resin was washed with 25 mM Tris pH 8.0, 320 mM KCl, 1 mM DTT and the PGL-3 (Δ425-452) molecules were eluted into 25 mM Tris pH 8.0, 500 mM KCl, 1 mM DTT. Finally, the eluate containing purified PGL-3 (Δ425-452)-6xHis-mEGFP was dialyzed into 25 mM HEPES pH 7.5, 1 M KCl, 1 mM DTT.

*GLH-1*: Insect cells expressing GLH-1-6xHis-mEGFP were resuspended in lysis buffer (25 mM HEPES pH 7.5, 300 mM KCl, 10 mM Imidazole, 1 mM DTT added with protease inhibitor cocktail Complete EDTA-free (Roche) and lysed using sonicator. The lysates were centrifuged at 21,000 × g in a JS-13.1 rotor (Beckman-Coulter), and Ni-NTA Agarose resin (Macherey-Nagel) was mixed with the supernatant to capture PGL-3 protein molecules. The resin was then washed with 25 mM HEPES pH 7.5, 300 mM KCl, 20 mM Imidazole. Finally, the GLH-1 molecules were eluted into 25 mM HEPES pH 7.5, 300 mM KCl, 250 mM Imidazole.

In preparation for anion-exchange chromatography, the eluate was diluted 1:6 in 25 mM Tris pH 8.0, 1 mM DTT and 4 ml of Q Sepharose resin (GE Healthcare) was mixed to capture GLH-1 protein molecules. After incubation for ~30 min, the resin was washed with 25 mM Tris pH 8.0, 1 mM DTT buffers containing 200 mM, 250 mM, 300 mM, 350 mM, 400 mM, 500 mM and 1 M KCl. The fractions with 250 to 350 mM KCl were pooled.

Pooled fractions were further purified using size exclusion chromatography with a HiLoad 16/60 Superdex 200 column (GE Healthcare) in 25 mM HEPES pH 7.5, 300 mM KCl, 1 mM DTT.

*Untagged PGL-3, S1, PGL-3 (1-452), PGL-3 (370-693), PGL-3 (K126E K129E), S1 (K126E K129E), PGL-1 and GLH-1* were purified by cleaving off 6xHis and mEGFP tags from the C-terminus of the recombinant proteins using 6xHis-tagged TEV protease, followed by incubation with Ni-NTA Agarose resin to remove 6xHis-tagged mEGFP and TEV protease.

### Size exclusion chromatography and multi-angle light scattering

Molecular weight analysis of PGL-3 (1-452) was carried out using an OMNISEC multi-detector instrument from Malvern Panalytical. All measurements were performed in 25 mM HEPES, 300 mM KCl, 1 mM TCEP at 25°C. Prior to measurement, samples were centrifuged at 10 000 g for 10 min. 90 µl of sample at 0.5 mg/ml was injected onto a Superose 6 10/300 column (Cytiva). The eluate from the column was fed into a refractive index detector followed by a right-angle light scattering detector (640 nm). Data were analysed using the OMNISEC software (v11.10) using a refractive index increment (dn/dc) of 0.185 ml/g. Bovine serum albumin (PierceTM) was used to calibrate the detectors. Samples were measured in duplicate.

### Labeling of proteins with fluorescent dye

In order to generate fluorescently labeled S1, PGL-1 and GLH-1, the proteins were first dialyzed into buffer containing 1 mM TCEP and then incubated with AlexaFluor 594 C5-maleimide (Invitrogen) in 25 mM HEPES pH 7.5, 300 mM KCl (1 M KCl for S1), 1 mM TCEP for 2 h at room temperature. To remove unbound dye, repeated buffer exchange was performed with 25 mM HEPES pH 7.5, 300 mM KCl (1 M KCl for S1), 1 mM TCEP until no fluorescence was detected in the flow through.

### Preparation of mRNA used in experiments

In order to track the localization of mRNA in our assays, fluorescently labeled mRNA was generated using in vitro transcription. Coding sequence of the act-4 gene from *C. elegans* was cloned to be transcribed from a T7 promoter. To generate in vitro transcribed act-4 mRNA, T7 RNA polymerase (New England BioLabs) was used and ribonucleotide solution mix was doped with 30% Aminoallyl-UTP-Cy5 (Jena Bioscience). The reaction was incubated for 1 h at 37 °C, then treated with DNase II and mRNA was finally purified using RNeasy Mini Kit (Qiagen). For all the other experiments, total mouse liver mRNA (Takara) was used.

### Imaging and fluorescence recovery after photobleaching (FRAP) experiments

Phase separation properties of PGL-3 constructs under various conditions were imaged within chambers created by attaching cover glass to glass slide with imaging spacer between them (Grace Bio-Labs SecureSeal, Sigma Aldrich). 4 μL of solution was placed at the center of the imaging chamber, such that the spacer had no contact with the solution. PGL-3 (1-452) (Fig. 1b) was imaged directly on the cover glass (without glass slide), as condensates of PGL-3 (1-452) were not stable when imaged within the glass chamber. Addition of 1% crowding agent PEG-35000 stabilized condensates of this N-terminal fragment of PGL-3 allowing imaging over time (Fig. 1c-e, 2a, 2b, Supplementary Fig. 2d). The surfaces of glass slide and cover glass were passivated through PEG-silanization as described earlier (Alberti et al., 2018).

The images were acquired with an inverted microscope (Olympus IX83) equipped with a spinning-disk scan head (4,000 rpm, Yokogawa CSU-W1), a sCMOS camera (pixel size 11um, Prime 95B Teledyne Photometrics), and a 100x/1.4 oil-immersion objective (Olympus UPLSAPO). For imaging and FRAP experiments, we used Coherent OBIS lasers of wavelengths 405nm (FRAP), 488 nm, 561 nm, and 640 nm. Single confocal plane was imaged over time for all FRAP experiments.

Phase separation properties of the PGL-3 constructs PGL-3 (1-577), PGL-3 (Δ425-452) and FRAP experiments of preheated PGL-3-mEGFP and S1-mEGFP were imaged within flow chambers created by attaching cover glass to glass slide with two double-sided tapes (~60 μm thick) positioned parallel to each other. The surface of the cover glass was passivated through PEG-silanization or coating with lipids as described earlier (Alberti et al., 2018). An inverted spinning-disk confocal imaging system was used, fitted with diode-pumped solid-state LASERs (wavelength 488 nm and 561 nm), Olympus UplanSApochromat 100x 1.4 NA oil-immersion objective, Yokogawa CSU-X1 (5000 rpm) spinning-disk scan head, and Andor iXon DU-897 BV back illuminated EMCCD camera.

### Image analysis

All image analysis was conducted using the Fiji image-processing package (http://fiji.sc/Fiji). Images used for analysis of FRAP experiments were corrected for camera noise and uneven illumination error. Intensities of the bleached areas were corrected for bleaching and normalized to prebleached image. FRAP data was fitted to single exponential function using Matlab (Supplementary figure 5e). All fits had R^2^ value of 0.92 or higher.

FRAP data for the construct S1(K126E K129E) was acquired with the same spinning-disk confocal imaging system as all other constructs, but a different FRAP filter was used. To account for any difference in signal intensity resulting from the use of a different FRAP filter, we compared the fluorescence intensities of S1-mEGFP, PGL-3-mEGFP and a uniformly green fluorescent slide (Chroma Autofluorescent Plastic Slides) in images acquired using the two different FRAP filters. This analysis using three different samples all gave similar values of correction factor around 2.6. We used this correction factor to normalize the intensities of S1(K126E K129E) and plotted the results in Fig. 5c.

### Phase separation assays

For phase separation assays, different recombinantly-purified PGL-3 constructs were used, and phase separation was triggered by diluting salt (KCl) concentration to either 150 mM or 100 mM in 25 mM HEPES, pH 7.5, 1 mM DTT. Following phase separation, condensates of PGL-3 (1-452) dissolved rapidly when incubated within glass-based imaging chambers. Addition of 1% crowding agent PEG-35000 stabilized condensates of this N-terminal fragment of PGL-3 allowing imaging over time.

In some cases, PGL-3 constructs were preheated at higher temperatures (43.5 °C or 50 °C) for 10 min, followed by incubation at room temperature for 10 mins and phase separation via dilution of salt concentration. Incubation at higher temperatures was carried out in a temperature-controlled water bath or PCR thermocycler. Heating untagged wild-type PGL-3 and the mutant S1 protein at 50 °C or 43.5 °C respectively resulted in aggregation. In contrast, presence of mEGFP tag reduced the extent of heat-induced aggregation to a large extent. Therefore, we used mEGFP-tagged PGL-3 constructs in these assays. Following incubation at 43.5 °C or 50 °C, we centrifuged the samples (5 min at 20,000 g) to separate the aggregates from the pool of soluble protein in the supernatant. To account for loss of protein via aggregation, we remeasured the concentration of protein in the supernatant and used this pool of mEGFP-tagged PGL-3 constructs for our assays.

At physiological concentrations (approx. 1 μM), the PGL-3 protein is unable to phase separate into condensates in vitro (Saha *et al*, 2016). At these concentrations, mRNA promotes phase separation of PGL-3 (Saha *et al*, 2016). To assay for mRNA-dependence of condensate assembly, it is therefore essential to use physiological concentrations of wild-type or mutant PGL-3 proteins (Figure 2). However, these condensates are generally too small to assay rate of internal rearrangement of PGL-3 molecules within condensates using fluorescence recovery after photobleaching experiments. Therefore, to generate large condensates for measuring internal rearrangement of wild-type or mutant PGL-3 molecules, we primarily used higher concentrations of these proteins in vitro (binding to RNA is not essential for phase separation at these concentrations). However, to mimic the in vivo P granule phase as closely as possible, we generally added constituent proteins in proportion to their in vivo abundance (Saha *et al*, 2016) as described below.

In assays investigating phase separation of two or more components (i.e. PGL-3, S1, different PGL-3 constructs, PGL-1, GLH-1 or mRNA), the components were mixed with each other before phase separation was triggered by diluting the salt (KCl) to final concentration of 100 mM in 25 mM HEPES pH 7.5, 1 mM DTT. In experiments where we studied the correlation between concentration of S1 or PGL-3 in condensates and their half-time of FRAP recovery: 1) all tested mixtures contained the same concentration of S1 (5 μM) or PGL-3 (5 μM) and 5% of the protein pool of S1 or PGL-3 was mEGFP-tagged; 2) Image acquisition settings were the same for all the experimental groups: 50 ms exposure; 488-laser power set to 100%; bleaching power set to 100%.

We used fluorescently labeled PGL-3 constructs, PGL-1, GLH-1 or act-4 mRNA to check if they colocalize with S1 or PGL-3 in condensates. All components were mixed with each other before phase separation was triggered by diluting the salt (KCl) to final concentration of 100 mM in 25 mM HEPES pH 7.5, 1 mM DTT. In these experiments, 5% of the pool of PGL-3 or S1 proteins was fluorescently labeled. 100% of the other components in the mix were labeled with a fluorescent probe. Colocalization of different PGL-3 constructs with wild-type PGL-3 or S1 was tested by adding the different proteins in equal amounts (final concentration of each component was 5 μM). In experiments where colocalization of different P granule components with PGL-3 or S1 was assayed, concentration (or concentration ratio) of the different components were the same as in FRAP experiments (6.7 μM PGL-1, and 14 ng/ μL mRNA with 5 μM S1 or 5 μM PGL-3; 1 μM GLH-1 with 5 μM PGL-3; 2 μM GLH-1 with 10 μM S1).

We tested if phase separation of S1 is reversible. We triggered phase separation of S1 (20 µM) by diluting salt (KCl) concentration to 100 mM in 25 mM HEPES, pH 7.5, 1 mM DTT. Fluorescence microscopy confirmed presence of condensates of S1. An aliquot containing pre-formed condensates of S1 was mixed with buffer containing 1 M KCl. Here, we did not detect condensates or aggregates of the S1 protein. Following addition of 1 M KCl containing buffer, the final conditions were 6,7 μM S1 in 25 mM HEPES pH 7.5, 700 mM KCl, 1 mM DTT.

### *C. elegans* strains, maintenance and transgenic strain construction

*C. elegans* strains were maintained on plates containing nematode growth media seeded with OP50 bacteria using standard methods (Brenner, 1974). The strains TH561, TH623 and SSA130 were generated using CRISPR/Cas9-based technology (Table 1). Strains were verified by sequencing of the modified locus.

**Table 1.**
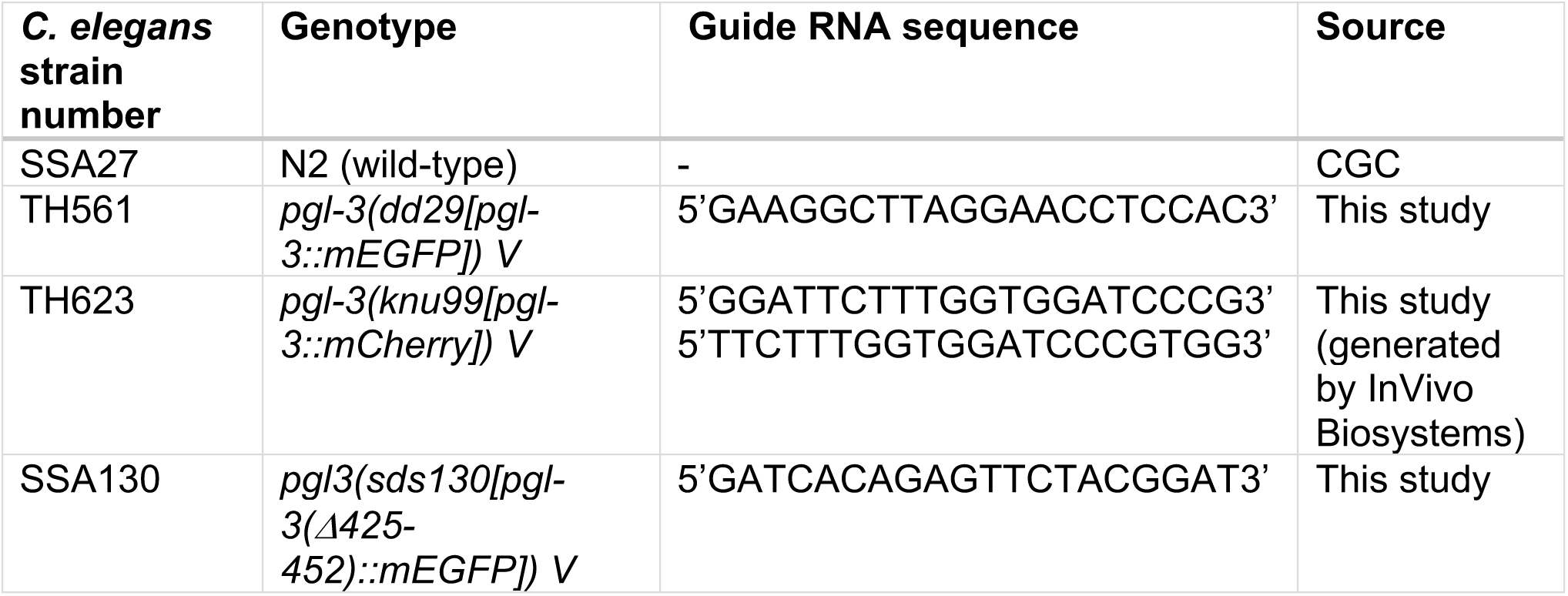
*C. elegans* strains used in this study.

### Fecundity assay

We assayed fecundity of the worms TH561 (*pgl-3::mEGFP*) and SS130 (*pgl-3(Δ425-452)::mEGFP*). These animals were grown and maintained at 20 °C throughout the experiment. L4 worms were singled for each strain. Each of these worms was transferred to a fresh plate every 24 h for five days. On the fourth day after transfer to a fresh plate, the number of progenies from each animal was scored. The total number of progenies for each worm was calculated and shown in Fig. 4d.

### Microinjection of recombinant proteins into *C. elegans* gonads

Recombinant PGL-3 constructs were microinjected into gonads of young adult *C. elegans* to study 1) their colocalization with P granules, and 2) the effect of the P granule phase on their dynamics. To study colocalization, mEGFP-tagged PGL-3 constructs were injected into gonads of worms expressing PGL-3-mCherry (strain TH623) from the native locus (to track P granules). Concentrations of injected proteins: PGL-3-mEGFP (10 μM), S1-mEGFP (6 μM) and PGL-3-mEGFP preheated at 50 °C (30 μM). To minimize the effect of fluorescent tag on dynamics, FRAP experiments were conducted on N2 worms microinjected with mEGFP-tagged PGL-3 constructs. Concentration of injected proteins: PGL-3-mEGFP (20 μM), S1-mEGFP (25 μM) and PGL-3-mEGFP preheated at 50 °C (30 μM). Worms were immobilized using 2 mM levamisole and imaged immediately following injections.

### Circular Dichroism (CD) spectroscopy

CD spectroscopy of PGL-3 constructs was carried out in a quartz cuvette (0.5 mm pathlength) using Chirascan Plus spectrometer (Applied Photophysics) equipped with TC1 temperature controller (Quantum Northwest). To measure the change in α-helical content of PGL-3 at different temperatures (data presented in Fig. 3b), the temperature of the cuvette was varied from 20 °C to 90 °C in 2 °C steps. At each temperature, the system was allowed to equilibrate for 180 s before measurement of CD signal between 200 and 260 nm. To test if unfolding of PGL-3 constructs at higher temperature is reversible once the temperature is lowered (data presented in Supplementary Fig. 3h and i), CD signal was measured initially at 21 °C, followed by measurements at a higher temperature (43.5 °C or 50 °C) and once again after the temperature was lowered back to 21 °C. At each temperature, the system was allowed to equilibrate for 15 min before measurement of CD signal between 200 and 260 nm.

### Statistical analysis

Two-tailed t test was performed for comparing two groups of data (number of condensates with or without addition of mRNA in Fig 2b; number of progeny in Fig 4d; estimated half time of recovery of fluorescence signal for bleached P granules in TH561 (*pgl-3::mEGFP*) and SSA130 (*pgl-3(Δ425-452)::mEGFP*) in Supplementary Fig. 4b; estimated half time of recovery of fluorescence signal for bleached condensates of S1 and PGL-3 with addition of PGL-1, GLH-1 and mRNA in Supplementary Fig. 4c). One way ANOVA was performed for comparing three groups of data (estimated half time of recovery of fluorescence signal for bleached P granules for injected proteins: PGL-3, PGL-3 preheated at 50 °C and S1 in Supplementary Fig. 4a). p values are indicated in the figure legends. Statistical tests were performed using GraphPad Prism version 8.4.3 (GraphPad Software, San Diego, California USA). To estimate the half time of recovery for above mentioned statistical analysis, the data points for normalized fluorescence signal for individual P granule/condensate were fit using R (RStudio Version 1.2.1335) to the following equation: F_t_ = (F_0_ + F_∞_*(t/t_1/2_))/(1 + t/t_1/2_) where F_t_ is fluorescence signal at time t, F_0_ is fluorescence signal immediately after photobleaching (minimal fluorescence signal), F_∞_ is maximal recovered fluorescence signal and t_1/2_ is half time of fluorescence recovery (Yguerabide *et al*, 1982). Only high-confidence fits [i.e., goodness of fit (p < 1E-11)] were used for statistical tests.

## Acknowledgments

We thank A.A. Hyman, R.V. Pappu, A. Goloborodko for feedback on the manuscript and helpful discussions. We thank A.A. Hyman and C. Hoege for the *C. elegans* strains TH623 and TH561. We thank Jana Sipkova, Anoop Kumar Yadav and Felicitas Arlt for help with image analysis and technical support. S.S. initiated this study as a postdoctoral fellow in the lab of A.A. Hyman. We thank the services and facilities at Vienna BioCenter for support - IMBA BioOptics Facility for support with imaging and image analysis; IMBA Molecular Biology Service for providing reagents for cloning and recombinant protein purification, and sequencing of constructs; VBCF Protein Technologies for help with recombinant protein expression and performing SEC-MALS experiment; IMBA Protein Chemistry Facility for verification of recombinant proteins using mass spectrometry; VBCF Computational Biology Training (A. Aszódi) for help with generating scripts for curve fitting. N2 strain was provided by the CGC, which is funded by NIH Office of Research Infrastructure Programs (P40 OD010440). S.J., J.B., P.C., B.R.P., and S.S. are funded by the Austrian Academy of Sciences. A.S.H. is scientific advisor at Dewpoint Therapeutics.

## Author contribution

Conceptualization, S.J. and S.S.; Methodology, S.J., A.S.H. and S.S.; Investigation, S.J., J.B., P.C., B.R.P., M.R., C.H., S.S.; Writing – Original Draft, S.J. and S.S.; Writing – Review & Editing, S.J., A.S.H., and S.S.; Funding Acquisition, S.S.; Resources, A.S.H. and S.S.; Supervision, S.S.

## Supplementary Information

**Supplementary Figure 1.**
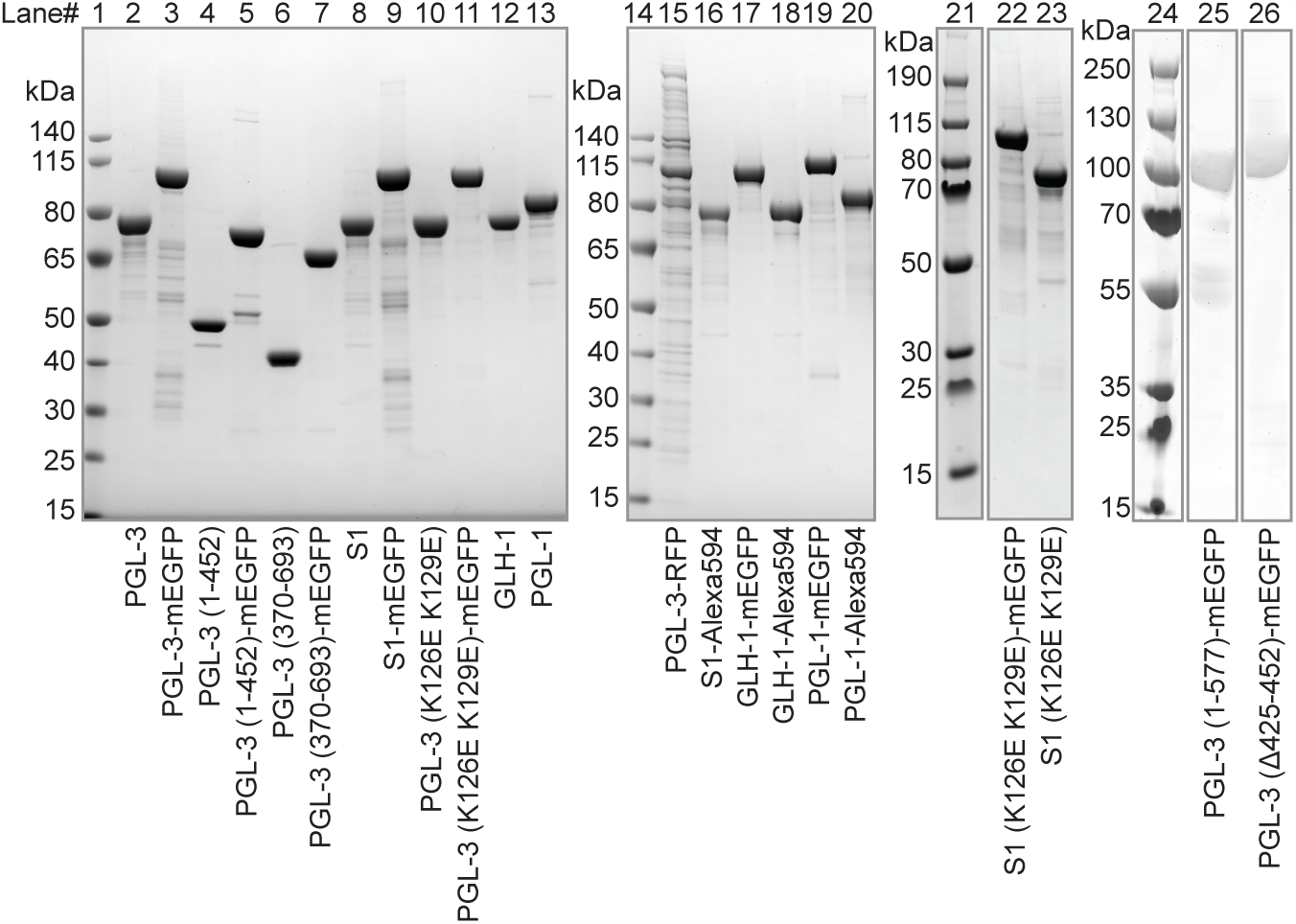
SDS-PAGE of recombinantly-purified proteins used in this study. Four SDS-PAGE gels are shown (gel 1: lanes 1-13; gel 2: lanes 14-20; gel 3: lanes 21-23, gel 4: lanes 24-26). Lane 21 is not adjacent to lanes 22 and 23 in gel 3; lanes 24, 25 and 26 represent non-adjacent lanes in gel 4.

**Supplementary Figure 2.**
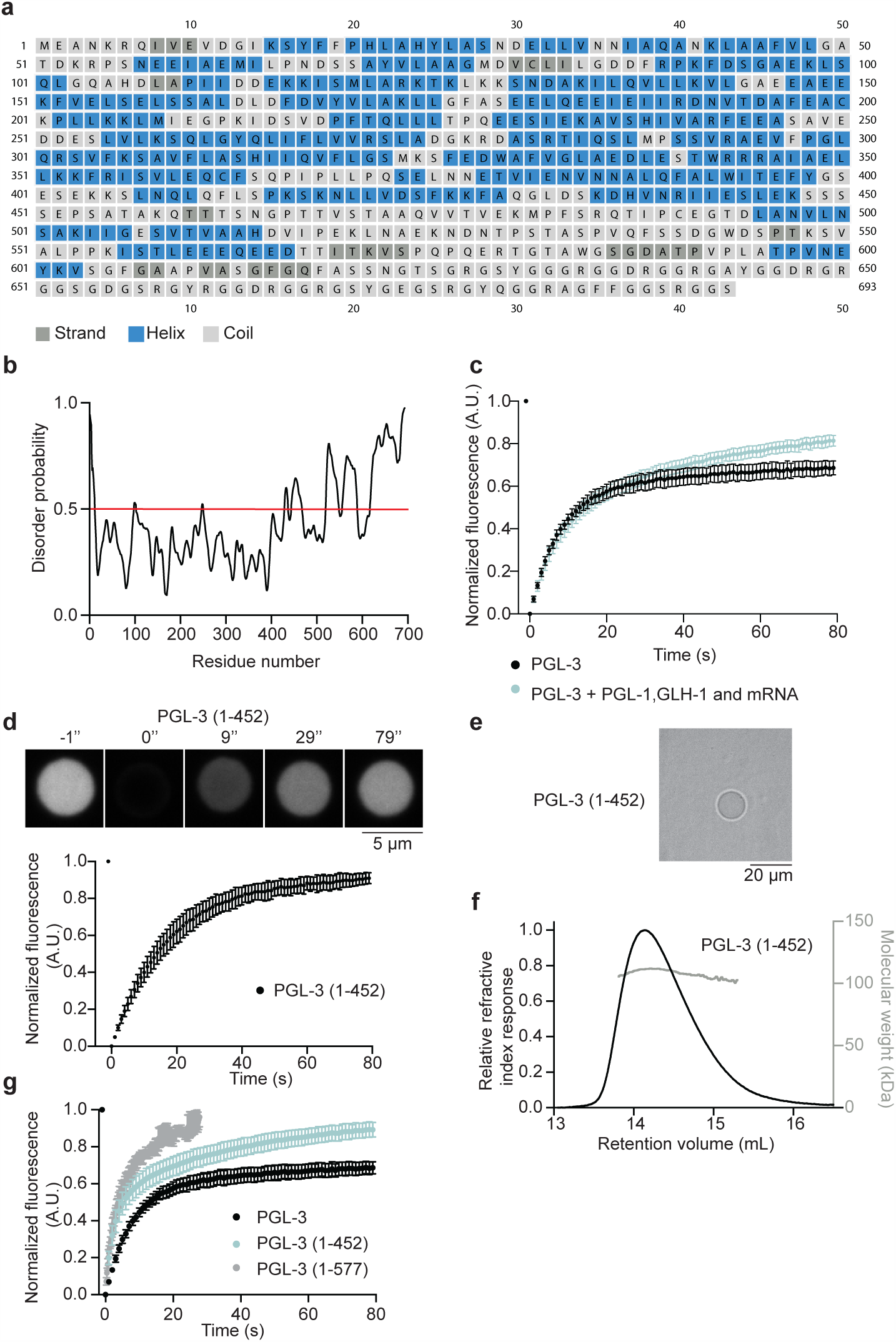
**a** Sequence of PGL-3 with secondary structure highlighted in different colors as predicted by the web-based PSIPRED 4.0 server. Dark grey ∞ strand, blue: α helix, light grey: coil. (http://bioinf.cs.ucl.ac.uk/psipred/) **b** Plot of disorder probability across the length of PGL-3, as predicted by the web-based PrDOS (Protein DisOrder prediction System) server with false-positive rate of 5.0% (http://bioinf.cs.ucl.ac.uk/psipred/). **c** Plot of average normalized fluorescence over time in bleached regions (1.5 μm × 1.5 μm) within condensates of PGL-3 alone (black circles, 10 μM, 5% mEGFP-tagged, n = 18), or PGL-3 (blue circles, 5 μM, 5% mEGFP-tagged, n = 21) in presence of PGL-1 (6.7 μM), GLH-1 (1 μM) and total mRNA (14 ng/μL). Error bars represent 1 SD. **d** Time-lapse fluorescence micrographs show fluorescence recovery over time after a condensate of PGL-3 (1-452) (5 μM, 5% mEGFP-tagged) was photobleached. Scale bar 5 μm. Below is shown a plot of average normalized fluorescence over time in 11 condensates of PGL-3 (1-452) (21 μM, 5% mEGFP-tagged) that were photobleached. Error bars represent 1 SD. **e** Bright field micrograph of PGL-3 (1-452) (5 μM) in 25 mM HEPES pH 7.5, 100 mM KCl, 1 mM DTT. Scale bar 20 μm. **f** Representative plot of size exclusion chromatography and multi-angle light scattering (SEC-MALS) data for PGL-3 (1-452). Black: relative refractive index response (Y-axis, left), Grey: molecular weight in kDa (Y-axis, right). Measurements were performed in 25 mM HEPES, 300 mM KCl, 1 mM TCEP. Average molecular weight of PGL-3 (1-452) estimated from SEC-MALS is 109.5 (±1.4) kDa (n = 2). Expected molecular weight based on amino acid sequence is 50.5 kDa, suggesting that PGL-3 (1-452) exists as a dimer. **g** Plot of average normalized fluorescence over time in bleached regions (1.5 μm × 1.5 μm) within condensates of PGL-3 (black circles, 5% mEGFP-tagged, n = 18), PGL-3 (1-452) (blue circles, 5% mEGFP-tagged, n = 24) and PGL-3 (1-577)-mEGFP (grey circles, bleached region 1.4 μm × 1.4 μm, n = 8). Error bars represent 1 SD.

**Supplementary Figure 3.**
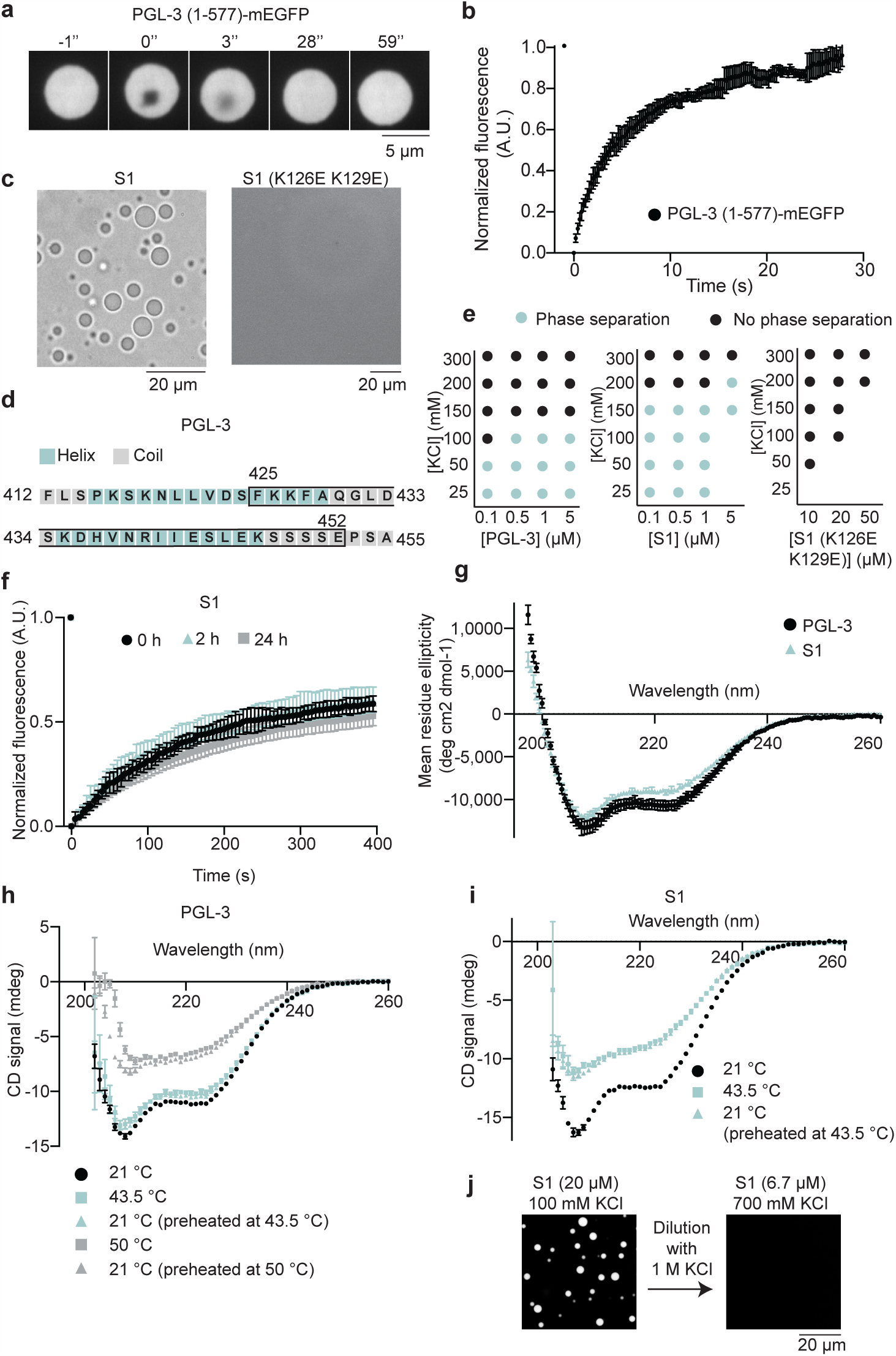
**a** Time-lapse fluorescence micrographs show fluorescence recovery over time after a 1.4 μm × 1.4 μm area within a condensate of PGL-3 (1-577)-mEGFP (20 μM) was photobleached. Scale bar 5 μm. **b** Plot showing average normalized fluorescence of bleached region (1.4 μm × 1.4 μm) within PGL-3 (1-577)-mEGFP condensates (n=8) over time. Error bars represent 1 SD. **c** Bright-field micrographs of S1 (8 μM) in 25 mM HEPES pH 7.5, 100 mM KCl, 1 mM DTT and S1 (K126E K129E) (50 μM) in 25 mM HEPES pH 7.5, 150 mM KCl, 1 mM DTT. Scale bar 20 μm. **d** Sequence of PGL-3 (412-455) with secondary structure highlighted in different colors as predicted by the web-based PSIPRED 4.0 server. Light blue: α helix, grey: coil. (http://bioinf.cs.ucl.ac.uk/psipred/). Residues 425-452 (shuffled in S1) are highlighted within open black box. **e** Phase diagrams of PGL-3, S1 and S1(K126E K129E) at different protein and salt (KCl) concentrations in 25 mM HEPES pH 7.5, 1 mM DTT. Blue circles represent conditions where phase separation was observed. Black circles represent conditions where phase separation was absent. 5% of the pool of protein was tagged with mEGFP in each case. **f** Plot showing average normalized fluorescence of bleached regions (1.5 μm × 1.5 μm) within condensates of S1 (10 μM, 5% mEGFP-tagged) over time. Condensates were assayed at different times after assembly via phase separation and incubation at 20 °C (Black circles: immediately following assembly (n = 16), blue triangles: 2 h post-assembly (n = 14), grey squares: 24 h post-assembly (n = 11). Error bars represent 1 SD. **g** Plot of mean residue ellipticity of PGL-3 and S1 vs. wavelength assayed using circular dichroism (CD) spectroscopy. Black circles: PGL-3 (3 μM, 10 mM Potassium phosphate pH 7.5, 300 mM KCl, 1 mM TCEP), Blue triangles: S1 (3 μM, 10 mM Potassium phosphate pH 7.5, 300 mM KCl, 1 mM TCEP). Error bars represent 1 SD among six CD scans. **h-i** Plots of circular dichroism signal vs. wavelength of **h** PGL-3 (3 µM) and **i** S1 (3 µM) in 25 mM HEPES pH 7.5, 1 M KCl and 1 mM DTT at different temperatures. Error bars represent 1 SD among six CD scans. Black circles: 21 °C, blue squares: 43.5 °C, blue triangles: 21 °C (after preheating at 43.5 °C), grey squares: 50 °C, grey triangles: 21 °C (after preheating at 50 °C). **j** Fluorescence micrographs show increasing salt (KCl) concentration dissolve pre-formed condensates of the S1 protein. 5% of total pool of the S1 protein was tagged with mEGFP. Condensates of S1 protein (20 μM) were prepared in 25 mM HEPES pH 7.5, 100 mM KCl, 1 mM DTT. After dilution with buffer containing 1 M KCl, the final concentration of S1 was 6,7 μM in 25 mM HEPES pH 7.5, 700 mM KCl, 1 mM DTT. Scale bar 20 μm.

**Supplementary Figure 4.**
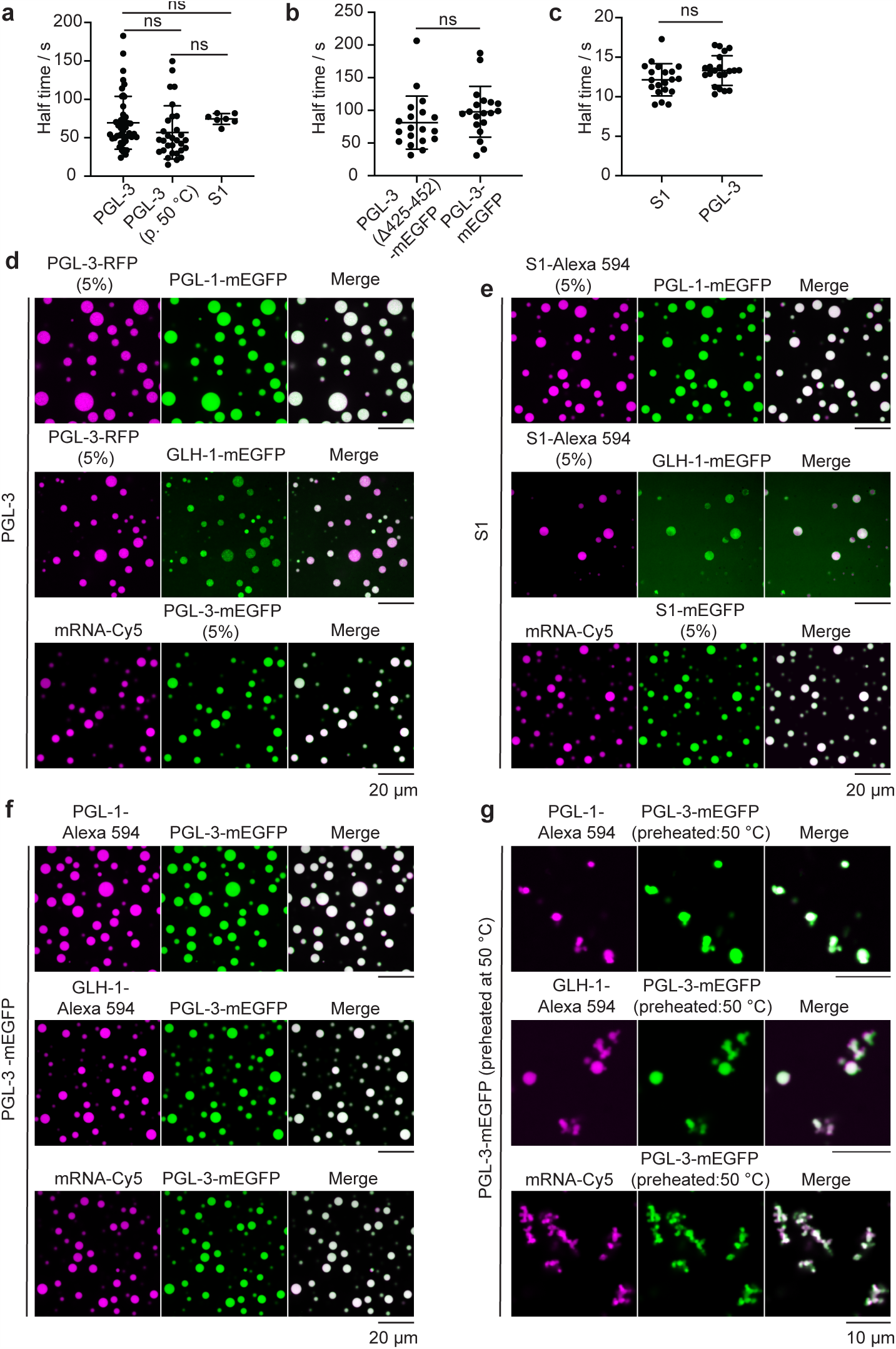
**a-c** Plots with estimated half times of recovery with statistical analysis for three experiments. Each black circle represents estimated half time (s) for one bleached P granule/condensate. The mean and 1 SD are also indicated. Half times were estimated by fitting the data points for normalized fluorescence signal for individual P granule/condensate to the equation: F_t_ = (F_0_ + F_∞_*(t/t_1/2_))/(1 + t/t_1/2)._ Only points with very confident fit (p<1E-11) were considered for the statistical analysis (described in more detail in Materials and Methods section). “ns” indicates no significant difference (p > 0.05) between the half times of recovery. Experiments were as following: **a** Bleaching of the P granules in live animals of N2 strain after PGL-3 (n = 46), preheated PGL-3 (n = 29) or S1 (n = 7) protein injection (Fig. 4b); One way ANOVA performed to compare the groups with following p values: p(PGL-3 vs. preheated PGL-3) = 0.241; p(PGL-3 vs. S1) = 0.926; p(preheated PGL-3 vs. S1) = 0.417; **b** Bleaching of the P granules in live animals in the strains expressing PGL-3 (Δ425-452)-mEGFP (n = 19) or PGL-3-mEGFP (n = 19) (Fig. 4b). T test performed to compare the two groups with resulting p value p = 0.205 **c** Bleaching of the condensates of PGL-3 (5 μM) (n=21) or S1 (5 μM) (n=20). 5% of the pool of protein was tagged with mEGFP) in presence of PGL-1 (6.7 μM), GLH-1 (1 μM), and total mRNA (14 ng/μL). T test performed to compare the two groups with resulting p value p = 0.104 **d-g** Fluorescence micrographs show concentration of PGL-1, GLH-1 and act-4 mRNA (in vitro transcribed) within condensates of PGL-3 **d**, S1 **e**, PGL-3-mGFP **f** and PGL-3-mEGFP preheated at 50 °C **g**. To track condensates of PGL-3 and S1 in **d-e**, 5% of the pool of PGL-3 or S1 proteins used were tagged with RFP, mEGFP or Alexa 594. Fluorophores used to track PGL-1 (mEGFP or Alexa 594), GLH-1 (mEGFP or Alexa 594), and mRNA (Cy5) are specified in each case. Scale bars **d-f** 20 μm, **g** 10 μm.

**Supplementary Figure 5.**
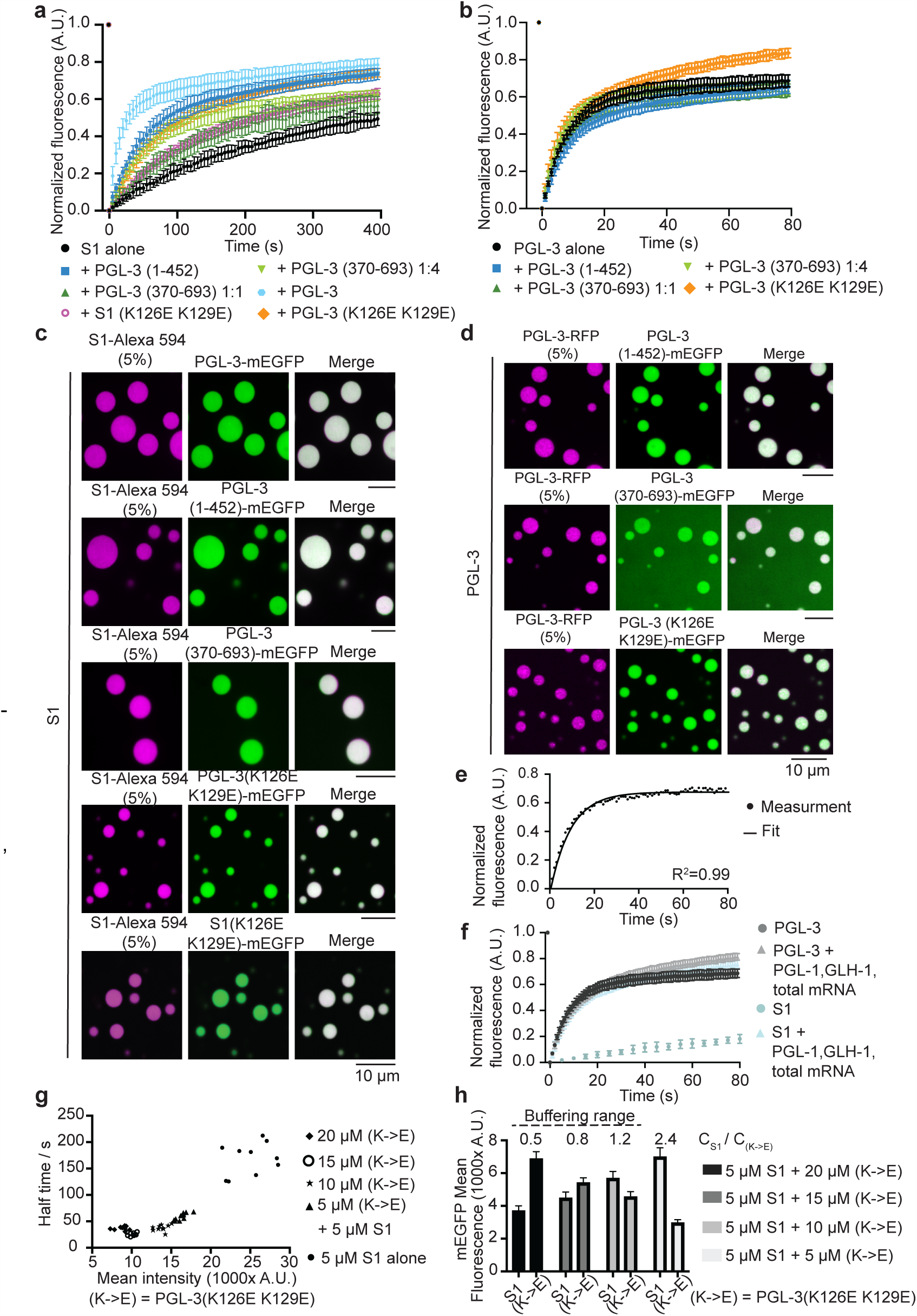
**a** Plot of average normalized S1-mEGFP fluorescence over time in bleached regions (1.5 μm × 1.5 μm) within the condensates of defined composition as follows. Black circles: S1 alone (10 μM, 5% mEGFP-tagged) (n = 18), light green triangle: S1 (5 μM, 5% mEGFP-tagged) + PGL-3 (370- 693) (20 μM) (n = 24), blue square: S1 (5 μM, 5% mEGFP-tagged) + PGL-3 (1-452) (5 μM) (n = 16), light blue hexagon: S1 (5 μM, 5% mEGFP-tagged) + PGL-3 (5 μM) (n = 20), green triangle: S1 (5 μM, 5% mEGFP-tagged) + PGL-3 (370-693) (5 μM) (n = 11), orange diamond: S1 (5 μM, 5% mEGFP-tagged) + PGL-3 (K126E K129E) (5 μM) (n = 17), magenta open circle: S1 (5 μM, 5% mEGFP-tagged) + S1 (K126E K129E) (5 μM) (n = 13). Error bars represent 1 SD. **b** Plot of average normalized PGL-3-mEGFP fluorescence over time μm) within the condensates of defined composition as follows. Black circles: PGL-3 alone (10 μM, 5% mEGFP-tagged) (n = 18), light green triangle: PGL-3 (5 μM, 5% mEGFP-tagged) + PGL-3 (370-693) (20 μM) (n = 12), blue square: PGL-3 (5 μM, 5% mEGFP-tagged) + PGL-3 (1-452) (5 μM) (n = 28), green triangle: PGL-3 (5 μM, 5% mEGFP-tagged) + PGL-3 (370-693) (5 μM) (n = 13), orange diamond: PGL-3 (5 μM, 5% mEGFP-tagged) + PGL-3 (K126E K129E) (5 μM) (n =12). Error bars represent 1 SD. **c-d** Fluorescence micrographs show concentration of PGL-3-mEGFP, PGL-3 (1-452)-mEGFP, PGL-3 (370-693)-mEGFP, PGL-3 (K126E K129E)-mEGFP or S1(K126E K129E)-mEGFP within condensates of S1 **c** or PGL-3 **d**. To track condensates of S1 and PGL-3 in **c-d**, 5% of the pool of S1 or PGL-3 proteins used were tagged with Alexa 594 or RFP. Scale bar 10 μm. **e** A representative plot of normalized PGL-3-mEGFP fluorescence over time in bleached regions (1.5 μm × 1.5 μm) within a condensate and corresponding single exponential fit. FRAP data was fitted to the single exponential function: y = a * (1 − exp(−b*x)). R^2^ values were ≥ 0.92 for all fits. **f** Plot of average normalized fluorescence over time in bleached regions (1.5 μm × 1.5 μm) within condensates of PGL-3 alone (black circles, 10 μM, 5% mEGFP-tagged, n = 18), PGL-3 in presence of PGL-1 (6.7 μM), GLH-1 (1 μM) and total mRNA (14 ng/μL) (grey triangles, 5 μM, 5% mEGFP-tagged, n = 21), S1 alone (blue circle, 10 μM, 5% mEGFP-tagged, n =18), or S1 in presence of PGL-1 (6.7 μM), GLH-1 (1 μM) and total mRNA (14 ng/μL) (light blue triangle, 5 μM, 5% mEGFP-tagged, n = 21). Error bars represent 1 SD. **g** Plot from Figure 5b shown here with additional data representing condensates with S1 alone. Plot of t_1/2_ vs. mean S1-mEGFP fluorescence intensity per unit area (before photobleaching) within condensates. Each point on the plot represents data from a single photobleached region within a condensate of defined composition which includes 5 μM S1 alone (solid circle) or 5 μM S1 and different concentrations of PGL-3 (K126E K129E) indicated as follows. Solid diamond: 20 μM PGL-3 (K126E K129E), open circle: 15 μM PGL-3 (K126E K129E), asterisk 10 μM PGL-3 (K126E K129E), solid triangle: 5 μM PGL-3 (K126E K129E). Ten points are plotted for each composition (nine for 20 μM PGL-3 (K126E K129E)). **h** Estimation of the ratio of intra-condensate concentrations of S1 and additives for conditions represented in Figure 5b. Plot showing average mEGFP fluorescence intensity per unit area in condensates containing S1 and PGL-3 (K126E K129E): from left to right in shades of gray of decreasing intensity represent condensates generated with 5 µM S1 and PGL-3 (K126E K129E) (20 μM or 15 μM or 10 μM or 5 μM). Different sets of condensates were used to estimate the intra-condensate concentration of S1 and PGL-3 (K126E K129E) separately, i.e. either 5% mEGFP-tagged S1 or 5% mEGFP-tagged PGL-3 (K126E K129E) was used. 38 condensates were used to estimate S1 or PGL-3 (K126E K129E) concentration in each case. Error bars represent 1 SD. Within the buffering range (highlighted using broken lines), t_1/2_ of S1 within condensates remains unchanged. Estimated ratio of intra-condensate concentrations of S1 and PGL-3 (K126E K129E) is specified above the plot.

**Supplementary Figure 6.**
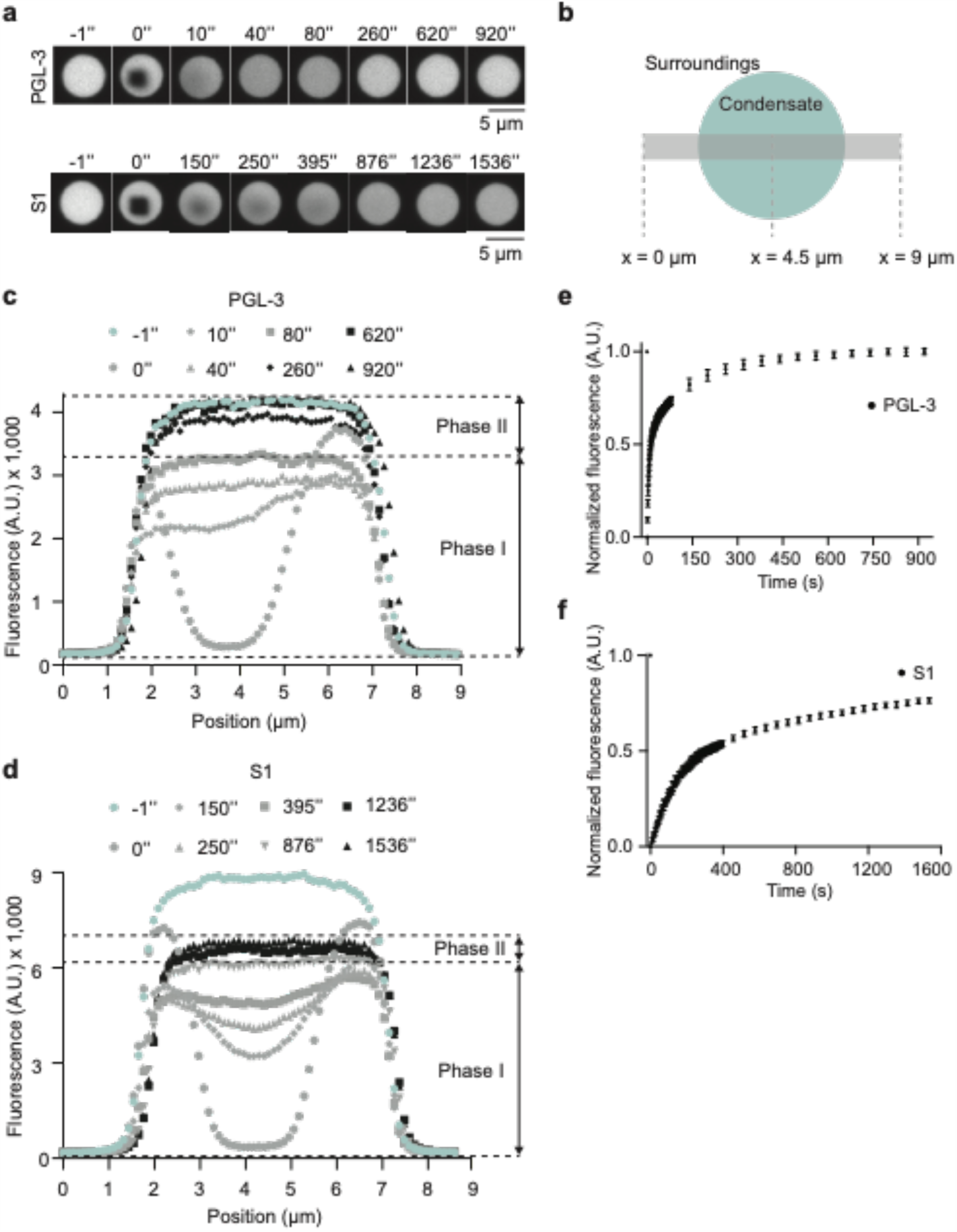
**a** Time-lapse fluorescence micrographs show fluorescence recovery after photobleaching over time in condensates of PGL-3 (10 μM) or S1 (10 μM) (5% of the pool of protein was tagged with mEGFP). Scale bars 5 μm. **b** Cartoon defining position with respect to a condensate. This position scheme was used to generate fluorescence-profile plots shown in **c** and **d**. **c-d** Profile plots mapping the fluorescence signal at different positions within a condensate of PGL-3 **c** or S1 **d**. Plots shows fluorescence signal in selected area (see **b** at different time points. Over time, fluorescence recovers within bleached regions through a combination of internal rearrangement of molecules within condensates and exchange of molecules between condensates and surroundings. Colors represent different recovery phases: blue for time before bleaching, grey for phase I and black for phase II. Phase I is a fast phase of fluorescence recovery that promotes homogenization of fluorescence signal within the condensate predominantly via internal rearrangement of molecules. Phase II is a slower phase of fluorescence recovery that depend predominantly on additional replenishment of unbleached molecules from the surroundings. Different time points are shown as follows. **c** Blue circle: −1 s (before bleaching), grey circle: 0 s (moment of bleaching), grey diamond: 10 s, grey triangle: 40 s, grey square: 80 s, black diamond: 260 s, black square: 620 s, black triangle: 920 s after bleaching. **d** Blue circle: −1 s (before bleaching), grey circle: 0 s (moment of bleaching), grey diamond: 150 s, grey triangle: 250 s, grey square: 395 s, grey reverse triangle: 876 s, black square: 1236 s, black triangle: 1536 s after bleaching. **e-f** Plots of average normalized fluorescence over time in bleached regions (1.5 μm × 1.5 μm) within the condensates of PGL-3 (n = 9) **e** or S1 (n=9) **f** imaged as described in **a**. Error bars represent 1 SD.

## Supplementary Movies

**Supplementary Movie 1**

Time-lapse fluorescence micrographs show fluorescence recovery after photobleaching of P granules in live *C. elegans* (pachytene region) expressing PGL-3-mEGFP. Time stamp and scale bar (3 μm) indicated (related to Fig. 4e).

**Supplementary Movie 2**

Time-lapse fluorescence micrographs show fluorescence recovery after photobleaching of P granules in live *C. elegans* (pachytene region) expressing PGL-3 (Δ425-452)-mEGFP. Time stamp and scale bar (3 μm) indicated (related to Fig. 4e).

